# The genetic and epigenetic landscape of the Arabidopsis centromeres

**DOI:** 10.1101/2021.05.30.446350

**Authors:** Matthew Naish, Michael Alonge, Piotr Wlodzimierz, Andrew J. Tock, Bradley W. Abramson, Christophe Lambing, Pallas Kuo, Natasha Yelina, Nolan Hartwick, Kelly Colt, Tetsuji Kakutani, Robert A. Martienssen, Alexandros Bousios, Todd P. Michael, Michael C. Schatz, Ian R. Henderson

## Abstract

Centromeres attach chromosomes to spindle microtubules during cell division and, despite this conserved role, show paradoxically rapid evolution and are typified by complex repeats. We used ultra-long-read sequencing to generate the Col-CEN *Arabidopsis thaliana* genome assembly that resolves all five centromeres. The centromeres consist of megabase-scale tandemly repeated satellite arrays, which support high CENH3 occupancy and are densely DNA methylated, with satellite variants private to each chromosome. CENH3 preferentially occupies satellites with least divergence and greatest higher-order repetition. The centromeres are invaded by *ATHILA* retrotransposons, which disrupt genetic and epigenetic organization of the centromeres. Crossover recombination is suppressed within the centromeres, yet low levels of meiotic DSBs occur that are regulated by DNA methylation. We propose that Arabidopsis centromeres are evolving via cycles of satellite homogenization and retrotransposon-driven diversification.

**One-sentence summary:** Long read sequencing and assembly of the Arabidopsis centromeres reveals their genetic and epigenetic topography.

## Introduction

Despite their conserved function during chromosome segregation, centromeres show diverse organization between species, ranging from single nucleosomes to megabase-scale tandem repeat arrays (*1*). Centromere ‘satellite’ repeat monomers are commonly ∼100–200 bp, with each repeat capable of hosting a CENPA/CENH3-variant nucleosome (*1, 2*). CENPA/CENH3 nucleosomes ultimately assemble the kinetochore and position spindle attachment on the chromosome, allowing segregation during cell division (*3*). Satellites are highly variable in sequence composition and length when compared between species (*2*). The library of centromere repeats present within a genome often shows concerted evolution, yet they have the capacity to change rapidly in structure and sequence within and between species (*1, 2, 4*). However, the genetic and epigenetic features that contribute to centromere evolution are incompletely understood, in large part due to the challenges of centromere sequence assembly and functional genomics of highly repetitive sequences.

*De novo* assembly of repetitive sequences is challenging. As such, most eukaryotic genome assemblies are in a fragmented state with many repetitive regions completely unresolved, especially the centromeres and other large repeats. Even the most scrutinized genomes, such as the human GRCh38 and Arabidopsis TAIR10 reference genomes, fail to represent centromeres and other large repeats of biological importance. However, recent advances in long-read sequencing, including Oxford Nanopore (ONT) and PacBio single-molecule technologies, have revolutionized the field by enabling substantially more complete and contiguous genome assemblies. Owing to their increased length and accuracy (10 kbp to >100 kbp with 90–99% mean accuracy), the long reads are capable of spanning and assembling repetitive sequences that are too ambiguous to resolve with previous sequencing technologies. Notably, using these technologies, the highly repetitive human centromeres have recently been assembled, leveraging the fact that sequence heterogeneity exists between the satellite repeats, effectively creating regularly spaced unique sequence markers (*5–10*). As such, given sufficiently long reads, a genome assembler can effectively bridge from one unique marker to the next, thereby creating a reliable and unambiguous reconstruction. This core concept, combined with more accurate base-calling and consensus generation, is now leading to highly accurate and complete representations of complex genomes for the first time (*5, 11*).

The *Arabidopsis thaliana* genome was first sequenced in 2000, yet the centromeres, telomeres, and ribosomal DNA repeats have remained unassembled, due to their high repetition and similarity (*12*). The Arabidopsis centromeres are known to contain millions of base pairs of the *CEN180* satellite repeat, which support CENH3 loading (*13–17*). We used ultra-long-read DNA sequencing to establish the Col-CEN reference assembly, which wholly resolves all five Arabidopsis centromeres. The assembly contains a library of 66,129 *CEN180* satellites, with each chromosome possessing largely private satellite variants. Higher-order *CEN180* repetition is prevalent within the centromeres and is also chromosome specific. We identify *ATHILA* LTR retrotransposons that have invaded the satellite arrays and interrupt centromere genetic and epigenetic organization. By analyzing functional data from mutant lines, we demonstrate that DNA methylation epigenetically silences initiation of meiotic DNA double-strand breaks (DSBs) within the centromeres. Together, our data are consistent with satellite homogenization and retrotransposon invasion driving cycles of centromere evolution in Arabidopsis.

### Complete assembly of the Arabidopsis centromeres

The *Arabidopsis thaliana* TAIR10 reference genome is an exceptionally accurate and complete eukaryotic assembly that is an invaluable resource for plant science (*12*). However, TAIR10 fails to represent the telomeres, some rDNAs and the centromere satellite arrays. To resolve these remaining sequences, we supplemented existing TAIR10 genomic resources with Oxford Nanopore (ONT) sequencing data from Columbia (Col-0) genomic DNA, comprising a total of 73.6 Gbp, and ∼55× coverage of ultra-long (>50 kbp) reads. This long-range sequence information, combined with our optimized assembly and validation pipeline, yielded a nearly closed and highly accurate assembly of the Col-0 genome (Col-CEN v1.0). Chromosomes 1 and 3 are wholly resolved from telomere-to-telomere (T2T), chromosomes 2 and 4 are complete apart from the *45S* clusters and adjacent telomeres, and a single gap remains on chromosome 5 (**Fig. 1**).

**Figure 1.**
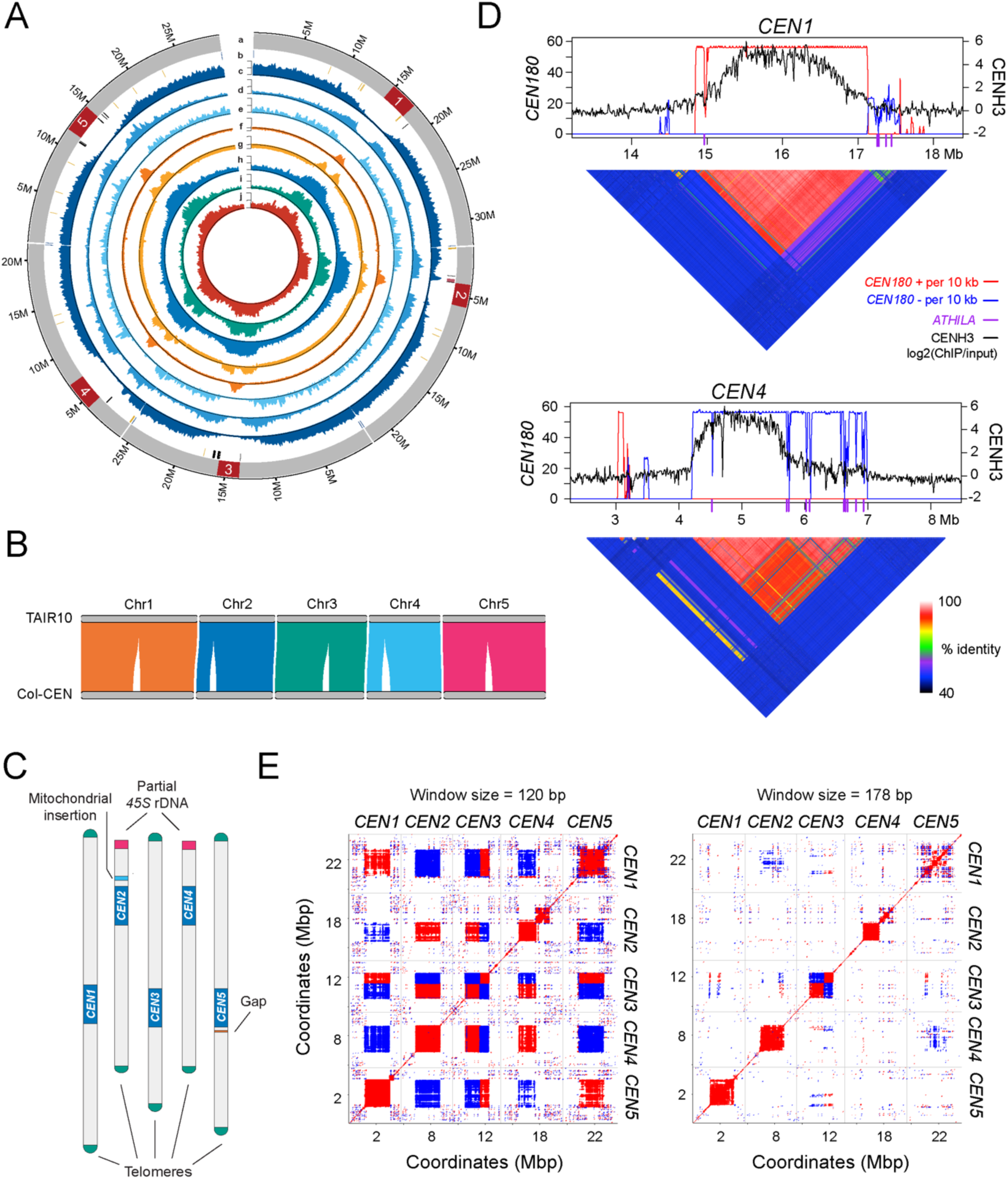
Complete assembly of the Arabidopsis centromeres. **A.** Genome-wide circos plot of the Col-CEN assembly. Quantitative tracks (c-j) are aggregated in 100 kbp bins and independent y-axis labels are given as (low tick value, mid tick value, high tick value, unit of measurement): (a) chromosome labels with centromeres shown in red; (b) genomic features showing telomeres in blue, *45S* rDNA in yellow, *5S* rDNA in black, and the Chr2 mitochondrial insertion in pink; (c) genes (0, 25, 51, # of genes); (d) transposable elements (0, 85, 171, # of transposable elements); (e) Col×Ler F_2_ crossovers (0, 6, 12, # of crossovers); (f) CENH3 (-1, 0, 3, log_2_(ChIP/Input)); (g) H3K9me2 (-1, 0, 2, log_2_(ChIP/Input)); (h) CpG methylation (0, 40, 80, % methylated); (i) CHG methylation (0, 20, 40, % methylated); (j) CHH methylation (0, 4, 8, % methylated). **B.** Plot showing syntenic alignments between the TAIR10 and Col-CEN assemblies. **C.** Genome assembly ideogram with annotated chromosome landmarks (not drawn to scale). **D.** CENH3 log_2_(ChIP/Input) (black) plotted over centromeres 1 and 4 (*13*). *CEN180* density per 10 kbp is plotted for forward (red) or reverse (blue) strand orientations. *ATHILA* retrotransposons are indicated by purple ticks on the x-axis. Beneath are heatmaps showing pairwise % sequence identity values of adjacent 5 kbp regions. **E.** Dotplot analysis comparing the 5 centromere regions, using a search window of 120 or 178 bp. Red and blue shading indicate detection of similarity on the same or opposite strands, respectively.

After repeat-aware polishing with R9 and R10 ONT reads and selective short-read polishing, the Col-CEN assembly is highly accurate with a QV of 33.95 and 45.58 inside and outside of the centromeres, equivalent to an error rate of 1 in 27,696 and 1 in 63,529 nucleotides, respectively (**Fig. S1A** and **Table S1**). The assembly is highly concordant with TAIR10, with 95.53% of Columbia BAC contigs aligning with high coverage and identity (>95%), and 99.61% of TAIR10 gene annotations represented in the assembly (97.49% of genes are exactly represented) (**Fig. 1B**). The Col-CEN assembly includes the *5S* rDNA arrays on chromosomes 3, 4, and 5, as well as a large mitochondrial genome insertion on chromosome 2 (**Fig. 1A and 1C**). Furthermore, the assembly reconstructs all five centromeres spanning 11,787,742 bp of new sequence, 120 and 98 kbp of *45S* rDNA in the chromosome 2 and 4 Nucleolar Organizing Regions (NORs), and the complete telomeres of the 8 chromosome arms without sub-telomeric NORs (**Fig. 1A–1C, S2 and S3**). We also identified a thionin gene cluster that was discordant with TAIR10 and after validating the structural accuracy of this locus in Col-CEN, we hypothesize that this may represent a TAIR10 misassembly, or a recent structural variant in our Col-0 line (**Fig. S3**).

The assembled centromere sequences are characterized by a repeated 178-bp motif (*CEN180*) that is organized into higher-order repeats (HORs) (**Fig. 1D, 2 and S4**). We validated the structural and base-level accuracy of the centromeres using techniques from the Human T2T consortium (*5*). Briefly, we aligned our Col-0 ONT reads to the assembly and observed even coverage across the centromeres, with few loci showing plausible alternate base signals (**Fig. S1**). We also observed relatively few ‘missing’ *k*-mers that are found in the assembly but not in Illumina short reads, which are diagnostic of residual consensus errors from the ONT reads (**Fig. S1**) (*18*). Notably, the five centromeres are relatively distinct at the sequence level, with each exhibiting chromosome-specific repeats (**Fig. 1E, 2 and Tables S2-S3**). This is consistent with our assembly pipeline unambiguously separating the five centromere sequences. We observe that unique ‘marker’ sequences are relatively frequent, with a maximum distance between consecutive markers in the assembled centromeres of only 28,630 bp, suggesting that our ultra-long reads can confidently span several unique markers and thus reliably assemble centromeric loci (**Fig. S1**).

**Figure 2.**
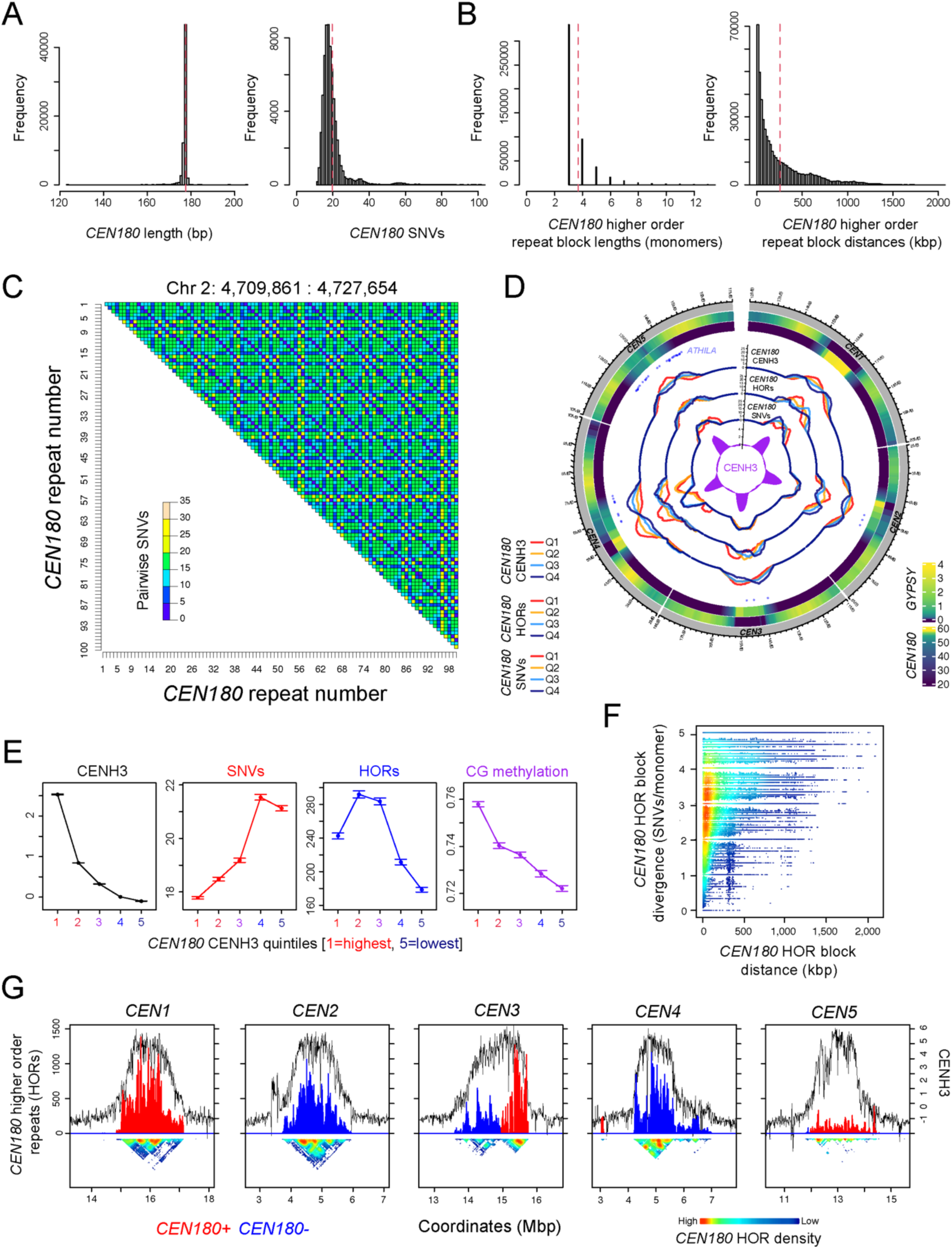
The Arabidopsis *CEN180* satellite repeat library. **A.** Histograms of *CEN180* monomer lengths (bp) and single nucleotide variants (SNVs) relative to the genome-wide consensus. Mean values are shown by the red dotted lines. **B.** As for A, but showing widths of *CEN180* higher order repeat (HOR) blocks (monomers), and the distance between HOR blocks (kbp). **C.** Heatmap of a representative satellite region within centromere 2, shaded according to pairwise SNVs between *CEN180*. **D.** Circos plot showing; (i) *GYPSY* LTR retrotransposon density, (ii) *CEN180* density, (iii) centromeric *ATHILA* rainfall plot, (iv) *CEN180* density grouped by decreasing CENH3 log_2_ (ChIP/Input) (red=high; navy=low), (v) *CEN180* density grouped by decreasing higher order repetition (red=high; navy=low), (vi) *CEN180* grouped by decreasing SNVs (red=high; navy=low) and, (vii) CENH3 log_2_(ChIP/Input), across the centromere regions. **E.** *CEN180* were divided into quintiles according to CENH3 log_2_(ChIP/Input) and mean values for each group with 95% confidence intervals plotted. The same groups were analyzed for *CEN180* SNVs (red), higher order repetition (blue) and CG context DNA methylation (purple). **F.** Plot of the distance between pairs of HOR blocks (kbp) and divergence (SNVs/monomers) between the HOR block sequences. **G.** Plots of CENH3 log_2_(ChIP/Input) (black) across the centromeres, compared to *CEN180* higher order repetition on forward (red) or reverse (blue) strands. A heat map is shown beneath that is shaded according to the density of *CEN180* higher order repeats.

### The Arabidopsis *CEN180* satellite repeat library

We performed *de novo* searches for tandem repeats to define the centromere satellite library (**Table S2**). We identified 66,129 *CEN180* satellites in total, with between 11,847 and 15,612 copies per chromosome (**Fig. 2 and Table S2**). The *CEN180* repeats form large stranded arrays, with the exception of centromere 3, which has an inverted structure (**Fig. 1D and S4**). The length of the repeat monomers is tightly constrained around 178 bp (**Fig. 2A**). We aligned all unique *CEN180* sequences (*n*=25,192) to derive a genome-wide satellite consensus. Each satellite was then compared to the consensus to calculate a single-nucleotide variant (SNV) score. Substantial sequence variation was observed between satellites, with a mean of 19.6 SNVs per *CEN180* **(Fig. 2A)**. Each centromere shows essentially private libraries of *CEN180* monomer sequences, with only 0.5% sharing an identical copy on a different chromosome (**Fig. 1E and Table S2**). In contrast, there is a high degree of *CEN180* repetition within chromosomes, with 54.2–65.4% showing one or more duplicates (**Table S2**). We also observed a minor class of ‘*CEN160*’ tandem repeats found mainly on chromosome 1 (1,289 repeats on Chr1, 43 repeats on Chr4, mean length=158.2 bp) (*17*).

We aligned CENH3 ChIP-seq data to the assembly and observed on average 10-fold log_2_(ChIP/Input) enrichment within the *CEN180* arrays, compared to the chromosome arms (**Fig. 1D and S4**) (*13*). CENH3 ChIP-seq enrichment is generally highest in the interior of the main *CEN180* arrays (**Fig. 1D and S4**). We observed a negative relationship between CENH3 ChIP-seq enrichment and *CEN180* SNV divergence (**Fig. 2D–2E**), consistent with CENH3 nucleosomes preferring to occupy satellites that are closer to the genome-wide consensus. In this respect, centromere 4 is noteworthy, as it consists of two distinct *CEN180* arrays, with the right array showing both higher SNV divergence and lower CENH3 ChIP-seq enrichment (**Fig. 1D, 2D and S4**). Together, this is consistent with satellite divergence leading to loss of CENH3 binding, or vice versa.

To define *CEN180* higher-order repeats (HORs), monomers were considered the same if they shared 5 or fewer pairwise SNVs. Consecutive repeats of at least 3 monomers below this SNV threshold were identified, yielding 500,833 HORs (**Fig. 2D and Table S3)**. Like the *CEN180* monomer sequences, HORs are almost exclusively chromosome specific (**Table S3**). The mean number of *CEN180* monomers per HOR was 3.69, equivalent to 656 bp (**Fig. 2B and Table S3**), and 91.1% of *CEN180* were part of at least one HOR. HOR block sizes show a negative exponential distribution, with the largest HOR formed of 60 monomers on chromosome 3, equivalent to 10,691 bp (**Fig. 2B**). Many HORs are in close proximity (42% are <100 kbp apart), although they are distributed along the length of the centromeres. For example, the average distance between HOR blocks was 250.7 kbp and the maximum distance was 2.1 Mbp **(Fig. 2B and Table S3)**. We also observed that HOR blocks that were a greater distance apart showed a higher level of SNVs between the blocks (SNVs/monomer) **(Fig. 2F)**, which is consistent with satellite homogenization being more effective over repeats that are physically closer.

The *CEN180* groups with highest CENH3 occupancy also show the greatest level of higher-order repetition and higher CG DNA methylation frequency **(Fig. 2D–2E and 2G)**. However, one notable exception to these trends is centromere 5, which harbours 12–22% of HORs compared to the other centromeres, yet still recruits comparable CENH3 **(Fig. 2G and Table S3)**.

### Invasion of the Arabidopsis centromeres by *ATHILA* retrotransposons

We observed that centromere 5 shows both reduced *CEN180* higher-order repetition and was heavily disrupted by breaks in the satellite array (**Fig. 2G and S4**). Genome-wide, within the main satellite arrays, the vast majority of sequence (>94%) is *CEN180*, with only 69 interspersed sequences larger than 1 kbp (**Table S4**). Within these gaps we identified 46 intact and 5 fragmented *ATHILA* LTR retrotransposons of the Gypsy superfamily, belonging to the *ATHILA*, *ATHILA2*, *ATHILA5* and *ATHILA6A/6B* subfamilies (**Fig. 3A and Table S4)** (*19–21*). The intact *ATHILA* elements have a mean length of 10.9 kbp, and the majority have highly-similar paired LTRs, target site duplications (TSDs), primer binding sites (PBS), polypurine tracts (PPT) and Gypsy superfamily open reading frames (**Table S4**). LTR comparisons indicate that the centromeric *ATHILA* elements are young, with on average 98.39% LTR sequence identity (**Fig. 3B and Table S4**), which was higher than *GYPSY* and *COPIA* elements located outside the centromere (**Fig. 3B**). We also observed 10 *ATHILA* solo LTRs that lacked a downstream PBS or upstream PPT, which is consistent with post-integration intra-element homologous recombination (**Table S4**). Interestingly, we also observed 5 instances where gaps containing full-length *ATHILA* or solo LTRs show a duplication on the same chromosome that are between 8.9 and 538.4 kbp apart, consistent with transposon sequences being copied post-integration, potentially via the same mechanism that generates *CEN180* HORs.

**Figure 3.**
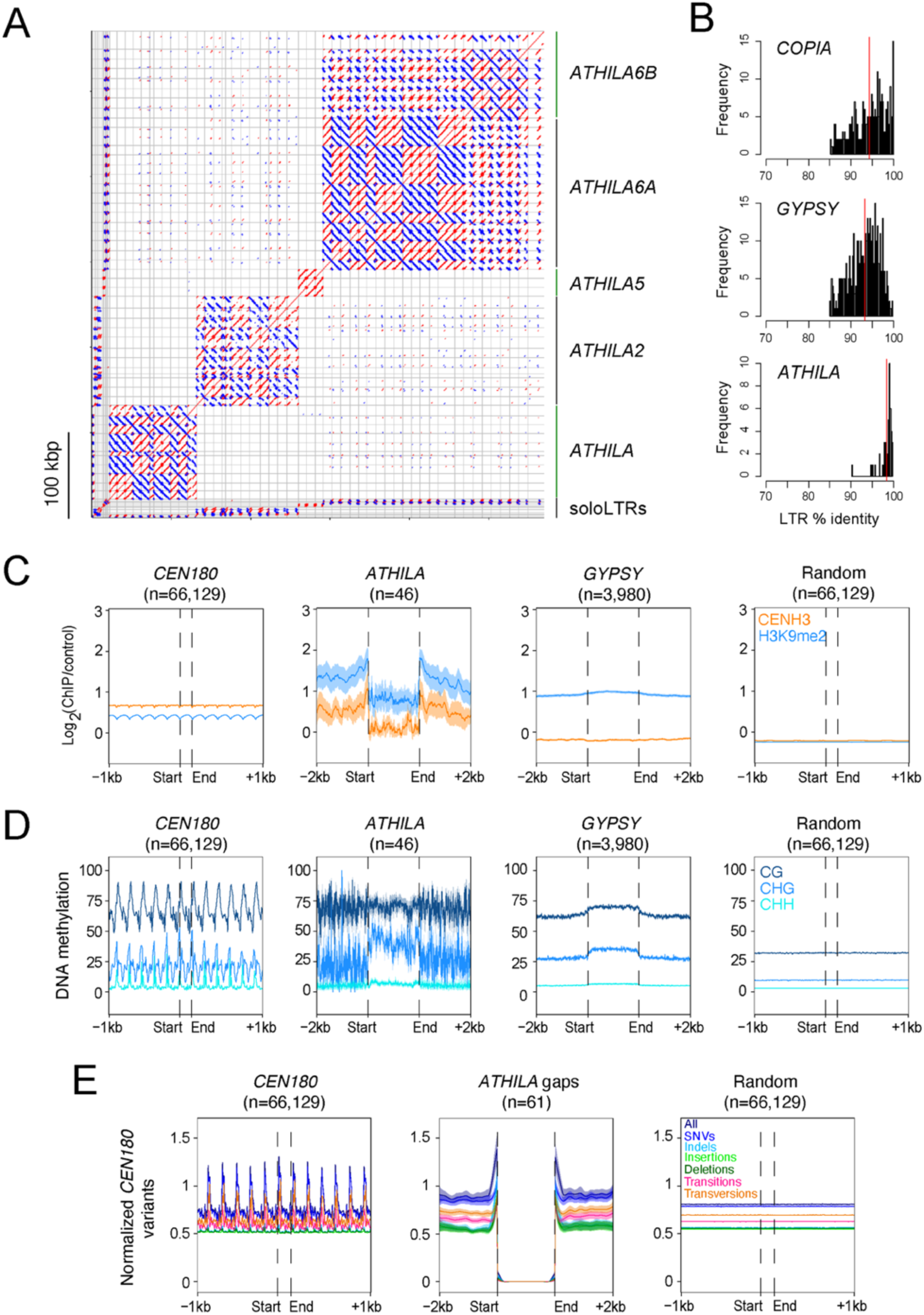
Invasion of the Arabidopsis centromeres by *ATHILA* retrotransposons. **A.** Dotplot of centromeric *ATHILA* retroelements using a search window of 50 bp. Red and blue indicate forward and reverse strand similarity. The elements assigned to different *ATHILA* families and solo LTRs are indicated. **B.** Histograms of LTR percent sequence identity for centromeric *ATHILA* elements, compared to *GYPSY* and *COPIA* elements outside of the centromeres. Mean values are indicated by the red lines. **C.** CENH3 (orange) and H3K9me2 (blue) log_2_(ChIP/Input) over *CEN180 (*n=66,129), centromeric intact *ATHILA (*n=46), *GYPSY* located outside the centromeres (n=3,980) and random positions (n=66,129). Shaded ribbons represent 95% confidence intervals for windowed mean values. **D.** As for C, but analyzing ONT-derived percent DNA methylation in CG (dark blue), CHG (blue) and CHH (light blue) contexts. **E.** The number of *CEN180* sequence edits (insertions, deletions, and mismatches, compared to the *CEN180* consensus) normalized by *CEN180* frequency, in positions surrounding *CEN180* (n=66,129), gaps containing *ATHILA* sequences (n=61), or random positions (n=66,129). All edits (dark blue) are analyzed, in addition to substitutions (SNVs, blue), indels (light blue), insertions (light green), deletions (dark green), transitions (pink) and transversions (orange).

We analyzed the centromeric *ATHILA* elements for CENH3 ChIP-seq enrichment and observed a decrease relative to the surrounding *CEN180*, yet higher levels than observed in *GYPSY* elements located outside the centromere (**Fig. 3C**). The *ATHILA* elements and flanking regions show greater H3K9me2 enrichment compared to *CEN180* (**Fig. 3C**). We used our ONT reads to profile DNA methylation over the *ATHILA* and observed dense methylation, at a similar level to the surrounding *CEN180*, although with higher CHG-context methylation (**Fig. 3D**). Hence, *ATHILA* elements are differentiated from the surrounding satellites at the chromatin level. Interestingly, when we profiled *CEN180* SNVs around gaps containing the *ATHILA* insertions (full length, fragments and solo LTRs), we observed a pronounced elevation in satellite divergence at the insertion boundaries (**Fig. 3E**). This may indicate that *ATHILA* integration was mutagenic on the surrounding satellite repeats, or that transposon insertion influenced the subsequent divergence or homogenization of the repeats. Together this indicates that centromeric *ATHILA* insertions interrupt the genetic and epigenetic organization of the Arabidopsis *CEN180* satellite arrays.

### Epigenetic organization and meiotic recombination within the centromeres

To assess genetic and epigenetic features of the centromeres, we analyzed all chromosome arms along their telomere–centromere axes using a proportional scale (**Fig. 4A**). Centromere midpoints were defined by maximum CENH3 ChIP-seq enrichment. As expected, *CEN180* satellites are highly enriched in proximity to the centromere midpoints (**Fig. 4A**). Gene density drops precipitously as the centromeres are approached, whereas transposons reciprocally increase, until they are replaced by *CEN180* (**Fig. 4A**). Gene and transposon density are tracked closely by H3K4me3 and H3K9me2 ChIP-seq enrichment, respectively (**Fig. 4A**). H3K9me2 enrichment is observed in the centromere, although there is a reduction in the centre coincident with CENH3 enrichment (**Fig. 4A**), consistent with reduced H3 occupancy caused by CENH3 replacement. Interestingly, a slight increase in H3K4me3 enrichment is observed within the centromeres, relative to the flanking pericentromeric regions (**Fig. 4A**). We observed striking biases in base composition over the centromeres, which are relatively GC-rich compared to the AT-rich chromosome arms (**Fig. 4A**).

**Figure 4.**
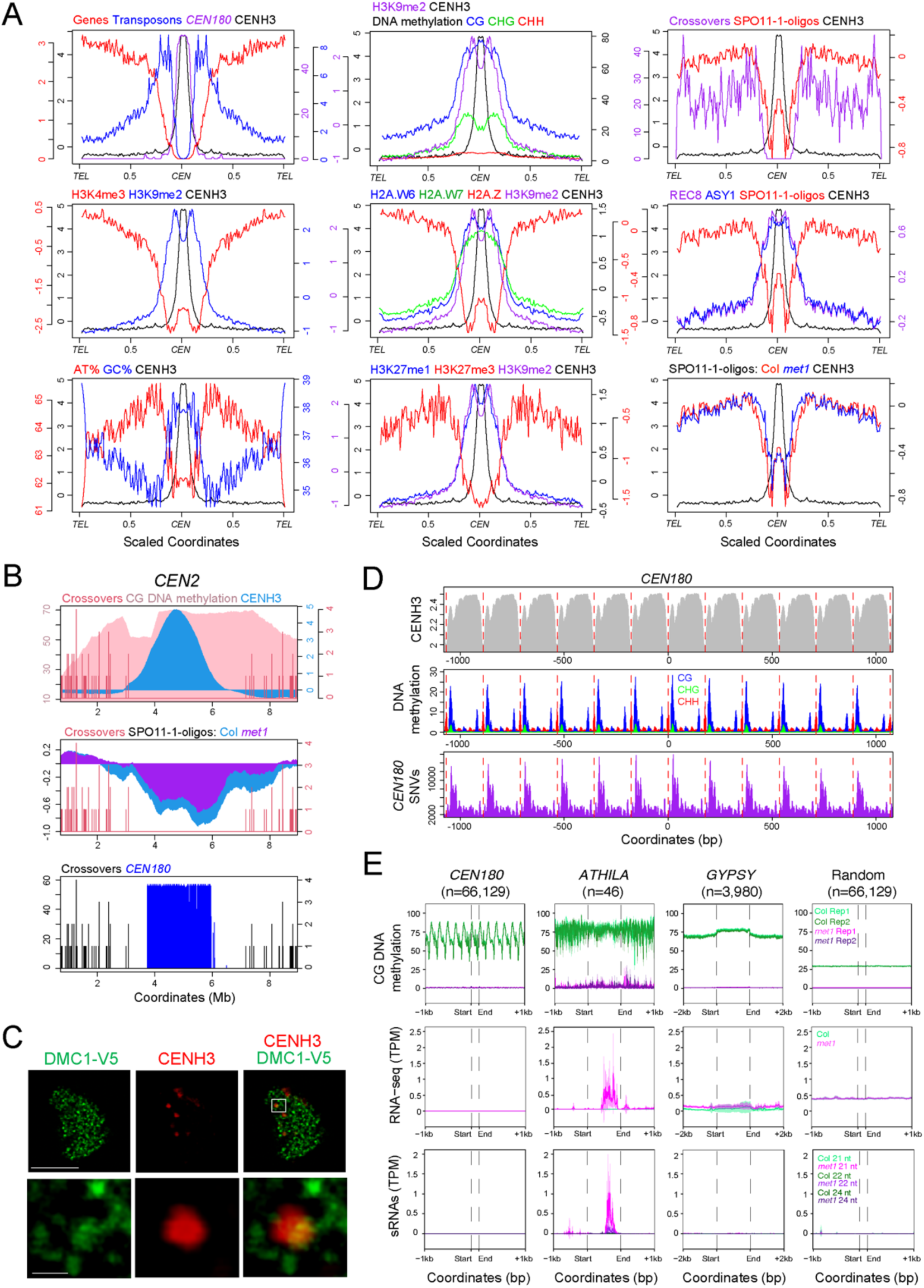
Epigenetic organization and meiotic recombination within the centromeres. **A.** Data were analyzed along chromosome arms that were proportionally scaled between the telomeres (*TEL*) and centromere midpoint (*CEN*), which was defined by maximum CENH3 log_2_(ChIP/Input). Data analyzed were gene, transposon and *CEN180* density, CENH3, H3K4me3, H3K9me2, H2A.W6, H2A.W7, H2A.Z, H3K27me1, H3K27me3, REC8 and ASY1 log_2_(ChIP/Input), % AT and GC base composition, DNA methylation, SPO11-1-oligos (in wild type and *met1*) and crossovers (see **Table S5** for information on datasets). **B.** Plot of crossovers (red/black), CG DNA methylation (pink), CENH3 (blue), SPO11-1-oligos in wild type (blue) and *met1* (purple) and *CEN180* density across centromere 2 (*CEN2*). **C.** Male meiocyte in early prophase I immunostained for CENH3 (red) and V5-DMC1 (green). Scale bars are 10 μM (upper row) and 1 μM (lower row). **D.** Plots of CENH3 ChIP enrichment (grey), % DNA methylation in CG (blue), CHG (green) and CHH (red) contexts and *CEN180* SNVs (purple), averaged over windows centred on all *CEN180* start coordinates. The red lines show 178 bp increments. **E.** CG context DNA methylation in wild type (green) or *met1* (purple) (*31*), RNA-seq in wild type (green) and *met1* (pink) (*30*), and siRNA-seq in wild type (green) and *met1* (pink) (*31*), over *CEN180* (n=66,129), centromeric intact *ATHILA (*n=46), *GYPSY* located outside the centromeres (n=3,980) and random positions (n=66,129). Shaded ribbons represent 95% confidence intervals for windowed mean values.

Using our ONT sequencing data and DeepSignal-plant, we observed dense DNA methylation across the centromeres in CG, CHG and CHH contexts (**Fig. 4A**) (*22*). However, CHG DNA methylation shows relatively reduced frequency within the centromeres, compared to CG methylation (**Fig. 4A**). This may reflect depletion of H3K9me2 within the centromeres, which functions to maintain DNA methylation in non-CG contexts (*23*). We also observed high centromeric ChIP-seq enrichment of the heterochromatic chromatin marks H2A.W6, H2A.W7 and H3K27me1 (**Fig. 4A**) (*24, 25*). The Polycomb-group modification H3K27me3 was depleted in the centromeres and found largely in the gene-rich chromosome arms (**Fig. 4A**). Enrichment of the euchromatic histone variant H2A.Z was low in the centromeres, but similar to H3K4me3, it showed a slight increase in the centromeres, relative to the pericentromeres (**Fig. 4A**). Hence, the centromeres show a unique chromatin state that is distinct from the euchromatic chromosome arms and heterochromatic pericentromeres.

To investigate genetic control of DNA methylation in the centromeres, we analyzed bisulfite sequencing (BS-seq) data from wild type and eight mutants defective in the CG and non-CG DNA methylation maintenance pathway (**Fig. S5**) (*23, 26*). Centromeric non-CG methylation is eliminated in a *drm1 drm2 cmt2 cmt3* mutant, and strongly reduced in *kyp suvh5 suvh6*, whereas CG methylation is intact in these lines (**Fig. S5**) (*23, 26*). CG methylation in the centromere is strongly reduced in *ddm1* and *met1*, although non-CG is more greatly reduced in *ddm1* than *met1* (**Fig. S5**) (*26*). Hence, dense DNA methylation is observed within the centromeres that is maintained by canonical pathways, although CG-context methylation is relatively high compared with non-CG.

Meiotic recombination, including unequal crossover and gene conversion, have been proposed to mediate centromere evolution (*4, 27*). We mapped 2,042 crossovers from Col×Ler F_2_ sequence data that were resolved on average to 1.01 kbp (**Fig. S6**). As expected, crossovers were potently suppressed in proximity to the centromeres (**Fig. 4A-4B and S6**). We observed high centromeric ChIP-seq enrichment of REC8-cohesin and the HORMA domain protein ASY1, which are components of the meiotic chromosome axis (**Fig. 4A**) (*28, 29*). To investigate the potential for meiotic DSB formation within the centromeres, we aligned SPO11-1-oligo data from wild type, which mark DSB sites (*30*). Overall, SPO11-1-oligos were low within the centromeres, although we observed an increase relative to the flanking pericentromeric heterochromatin, reminiscent of the H3K4me3 and H2A.Z patterns (**Fig. 4A**). To investigate the role of DNA methylation, we mapped SPO11-1-oligos sequenced in the CG DNA methylation mutant *met1-3* (*30*), which showed a gain of DSBs in proximity to the centromere (**Fig. 4A-4B**). To provide cytological evidence of recombination close to the centromeres, we immunostained meiocytes in early prophase I for CENH3 and V5-DMC1, which is a marker of inter-homolog recombination (**Fig. 4C and S7-S8)**. DMC1-V5 foci were observed along the chromosomes and associated with the surface, but not within, CENH3 foci (**Fig. 4C**). Hence, despite suppression of crossovers, we observe evidence for low levels of meiotic recombination initiation associated with the centromeres, which is influenced by DNA methylation.

Finally, we analyzed chromatin and transcription around *CEN180* and *ATHILA* retrotransposons at the fine-scale and compared wild type and the DNA methylation mutant *met1-3*. CENH3 nucleosomes show a strongly phased pattern of enrichment with the *CEN180* satellites, with relative depletion in spacer regions at the start and end of the satellites (**Fig. 4D**). Interestingly, these CENH3 spacer regions also associate with elevated DNA methylation and *CEN180* SNVs (**Fig. 4D**), consistent with CENH3-nucleosome occupancy influencing epigenetic modification and genetic divergence of satellites. In *met1*, we observed loss of CG-context DNA methylation in both the *ATHILA* and *CEN180* repeats (**Fig. 4E and S5**). However, RNA-seq and siRNA-seq counts increased specifically in the *ATHILA* in *met1* (**Fig. 4E**) (*30, 31*). Both RNA-seq and siRNA-seq signals increased most strongly in the internal 3′-regions of the *ATHILA* (**Fig. 4E**), which correspond to ‘TSI’ transcripts and easiRNA populations previously reported (*32–34*). This further indicates that epigenetic regulation of the *CEN180* satellites and *ATHILA* elements are distinct.

### Arabidopsis centromere evolution via satellite homogenization and *ATHILA* invasion

The Col-CEN assembly reveals the architecture of the Arabidopsis centromeres, which consist of megabase-scale, stranded *CEN180* arrays, which are invaded by *ATHILA* retrotransposons. Extensive sequence variation is observed between the satellites, and the majority of variant monomer sequences are private to each centromere. This is consistent with satellite homogenization occurring primarily within chromosomes. *CEN180* that are the least divergent and with greatest higher-order repetition show the highest CENH3 occupancy. This suggests that CENH3 chromatin may promote recombination pathways that lead to homogenization, including DSB formation and repair via homologous recombination. For example, inter-homolog strand invasion during meiosis has the potential to cause *CEN180* gene conversion. In this respect, we note that *CEN180* higher-order repeats show an average length of 656 bp, which is within the range of observed Arabidopsis meiotic gene conversions (*35*). We also see a proximity effect on divergence between higher-order repeats, with repeat blocks further apart showing greater sequence differences. These patterns are reminiscent of human alpha satellite higher-order repeats, although the alpha satellite blocks are longer and occur over greater distances (*5, 36, 37*). As meiotic crossover repair is strongly suppressed within the Arabidopsis centromeres, consistent with patterns across eukaryotes (*27, 38–40*), we do not consider unequal crossover to be likely within the centromeres. However, we propose that an ongoing, recombination-based homogenization process maintains the *CEN180* library close to the consensus that is optimal for CENH3 recruitment.

Aside from homogenizing recombination within the *CEN180*, the centromeres have experienced invasion by *ATHILA* retrotransposons. The ability of *ATHILA* elements to insert within Arabidopsis *CEN180* regions is likely determined by their integrase protein (*20, 41*). Interestingly, the Tal1 *COPIA* element from *Arabidopsis lyrata* shows a strong insertion bias into the *CEN180* when expressed in *Arabidopsis thaliana* (*42*), despite satellite sequences varying between these species (*43*). The majority of the centromeric *ATHILA* elements appear young, based on LTR identity and possess many features required for transposition, although the centromeres show striking differences in the frequency of *ATHILA* insertions, with centromeres 4 and 5 being the most invaded. *ATHILA* elements show lower CENH3 and higher H3K9me2 and CHG DNA methylation than the surrounding *CEN180*, and associate with increased satellite divergence at their boundaries. Hence, *ATHILA* represent a disruptive influence on the genetic and epigenetic organization of the centromeres. In maize, meiotic gene conversion was observed to act on *CRM2* retrotransposons within the centromeres (*27*). Therefore, satellite homogenization pathways may serve as a mechanism to eliminate *ATHILA* insertions. Indeed, a gene conversion mechanism may explain the five *ATHILA* intra-chromosome duplications that appear to have occurred post-integration. We also note that the presence of *ATHILA* solo LTRs is consistent with homologous recombination acting on the centromeric retrotransposons following integration. Intriguingly, centromere 5 and the diverged *CEN180* array on chromosome 4, show both high *ATHILA* density and a striking reduction of *CEN180* higher-order repetition. This is consistent with *ATHILA* inhibiting *CEN180* homogenization, or loss of homogenization facilitating *ATHILA* insertion, or both. We propose that each Arabidopsis centromere represents different stages in a cycle of satellite homogenization and disruption by *ATHILA*. These opposing forces provide both a capacity for homeostasis, and a capacity for change, during centromere evolution.

## Acknowledgements

We thank Isabel Thompson for *ATHILA* analysis. This work was supported by BBSRC grants BB/S006842/1 and BB/V003984/1, European Research Council Consolidator Award ERC-2015-CoG-681987 ‘SynthHotSpot’, Marie Curie International Training Network ‘MEICOM’ to IRH, US National Science Foundation grants DBI-1350041 and IOS-1732253 to MCS, and Royal Society awards UF160222 and RGF/R1/180006 to AB. The authors have no competing interests.

## Data availability

The ONT sequencing reads used for the Col-CEN assembly are available for download at ArrayExpress accession E-MTAB-10272 (http://www.ebi.ac.uk/arrayexpress/) (Username: Reviewer_E-MTAB-10272 Password: YVJAaVii). All data, code and materials are available in the manuscript or the supplementary materials.

## Author contributions

MN sequenced DNA, performed genome assembly, DNA methylation analysis and wrote the manuscript. MA performed genome assembly, polishing, validation, annotation and analysis and wrote the manuscript. PW performed satellite repeat annotation, genome analysis and wrote the manuscript. AJT performed short read alignment, genome analysis and wrote the manuscript. BA sequenced DNA and contributed to the assembly. CL performed meiotic immunocytology. PK performed meiotic immunocytology. NH sequenced DNA and contributed to the assembly. KC sequenced DNA and contributed to the assembly. TK provided intellectual input. RM provided intellectual input. AB performed *ATHILA* transposon annotation and genome analysis. TM supervised DNA sequencing, genome assembly and analysis. MS supervised genome assembly, validation, annotation and analysis and wrote the manuscript. IH supervised DNA sequencing, genome assembly, validation, annotation and analysis and wrote the manuscript.

## Supplementary Materials

### Materials and Methods

#### Genomic DNA extraction and ONT sequencing

For genomic DNA extraction, 3 week old Col-0 seedlings were grown on ½ MS media and 1% sucrose and kept in the dark for 48 hours prior to harvesting. Approximately 10 g of tissue was used per 200 ml of MPD-Based Extraction Buffer pH 6 (MEB). Tissue was flash frozen and the tissue ground in liquid nitrogen, using a pestle and mortar, and resuspended in 200 ml MEB. Ground tissue was thawed in MEB with frequent stirring. The homogenate was forced through 4 layers of miracloth, and then filtering again through 4 layers of fresh miracloth by gravity. 20% Triton x-100 was added to a final concentration of 0.5% on ice, followed by incubation with agitation on ice for 30 minutes. The suspension was centrifuged at 800*g* for 20 minutes at 4°C. The supernatant was removed and the pellet resuspended using a paintbrush in 10 ml 2-methyl-2,4 pentanediol buffer pH 7.0 (MPDB). The suspension was centrifuged at 650*g* for 20 minutes at 4°C. The supernatant was removed and the pellet was washed with 10 ml of MPDB. Washing and centrifugation was repeated until the pellet appeared white and was finally resuspended in a minimal volume of MPDB. From this point onwards all transfers were performed using wide bore pipette tips. 5 ml CTAB buffer was added to the nuclei pellet and mixed via gentle inversion, followed by incubation at 60°C until full lysis had occurred, taking between 30 minutes and 2 hours. An equal volume of chloroform was added and incubated on a rocking platform, with a speed of 18 cycles per minute, for 30 minutes, followed by centrifugation at 3000*g* for 10 minutes. An equal volume of phenol/chloroform/isoamyl alcohol (PCI, 25:24:1) was added to the lysate, followed by incubation on a rocking platform (18 cycles per minute) for 30 minutes. The lysate was centrifuged at 3000*g* for 10 minutes and the upper aqueous phase was transferred into a fresh tube. The PCI extraction was then repeated. The extraction was then repeated using only chloroform. 1/10^th^ volume of 3M Sodium Acetate was added to the lysate and mixed by gentle inversion. Two volumes of ice-cold ethanol were added and mixed by inversion. DNA was precipitated at -20°C for 48 hours. The precipitated DNA was removed using a glass hook and washed three times in fresh 70% ethanol. The DNA was dissolved in 120 µl of 10 mM Tris-Cl (pH 8.5).

Approximately 5 µg of DNA was size selected to be >30 kbp, using the BluePippin™ Size-Selection System (Sage Science) and the 0.75% DF Marker U1 cassette definition, with Range mode and BP start set at 30,000 bp. Library preparation followed the Nanopore SQK-LSK109 protocol and kit. Approximately 1.2-1.5 µg of size selected DNA in a volume of 48 µl was used for library preparation. DNA was nic-repaired and end-prepped by the addition of 3.5 μl of NEBNext FFPE Buffer and NEBNext Ultra II End Prep Reaction Buffer, followed by 2 µl of NEBNext DNA Repair Mix and 3 μl NEBNext Ultra II End Prep Enzyme Mix (New England Biolab, E7180S), with incubation for 30 minutes at 20°C, followed by 30 minutes at 65°C. The sample was cleaned using 1×volume AMPure XP beads and eluted in 61 μl of nuclease-free water. Adapters were ligated at room temperature using 25 µl Ligation Buffer, 10 µl NEBNext T4 DNA Ligase and 5 µl Adapter Mix for 2 hours. The library was cleaned with 0.4×volume AMPure XP beads, washed using ONT Long Fragment buffer and eluted in 15 µl elution buffer.

#### Col-CEN genome assembly

Libraries were sequenced on 6 ONT R9 flow cells and 1 ONT R10 flow cell, and the resulting .fast5 files were basecalled with Guppy (v4.0.15), using the dna_r9.4.1_450bps_hac.cfg and dna_r10.3_450bps_hac.cfg configurations, respectively. This yielded 73.6 Gb of sequence and ∼55x coverage of ultra-long reads (>50 kbp). The fastq files of ONT reads used for genome assembly are available for download at ArrayExpress accession E-MTAB-10272 (http://www.ebi.ac.uk/arrayexpress/). We trimmed adapters using Porechop (v0.2.4) and filtered for read lengths greater than 30 kbp and mean read quality scores >90%, using Filtlong (v0.2.0) (https://github.com/rrwick/Filtlong), which yielded 436,146 reads with a mean length of 43.9 kbp (19.15 Gbp), equivalent to 161× coverage of the TAIR10 genome. Flye (version 2.7) was used to assemble the reads, specifying a minimum read overlap of 10 kbp and a *k*-mer size of 17 (*44*).

#### Contig screen

We performed a comprehensive contig screen using methods inspired by the Vertebrate Genomes Project (VGP), though adapted for an inbred plant genome (*45*). We first aligned Flye contigs to the Columbia reference chloroplast (GenBank accession NC_000932.1) (*46*), and mitochondria (GenBank accession NC_037304.1) (*47*) genomes with Minimap2 (v2.17-r941, -x asm5) (*48*). Contigs with at least 50% of their bases covered by alignments were considered to be chloroplast or mitochondria genome sequences and were removed from the assembly.

We next used BLAST to screen for contigs representing bacterial contamination. We first masked the Flye assembly with windowmasker (v1.0.0, -mk_counts -genome_size 131405362) (*49*). We then aligned the Flye contigs to all RefSeq bacterial genomes (downloaded on 2020/05/21) with megablast (v2.5.0, -outfmt “6 std score”), providing the windowmasker annotations with “-window_masker_db” (*50*). We removed BLAST alignments with an E value greater than or equal to 0.0001, a score less than 500, and a percent identity less than 98%, and any contigs (four in total) with remaining alignments were manually inspected. Two of the four contigs were already identified as being chloroplast or mitochondria sequence and the other two were clearly nuclear contigs, so we determined that no contigs were derived from bacterial contaminants.

After removing chloroplast and mitochondria contigs, we performed one final screen to remove contigs with low read support. We aligned ONT reads (>=40 kbp) to the contigs with Minimap2 (v2.17-r941, - x map-ont) and removed any contigs (one in total) with more than 50% of its bases covered by fewer than 15 reads. Though we did not use its standard pipeline, we made use of purge_dups scripts for this analysis (*51*). After screening, the assembly consisted of 10 contigs with an N50 of 22,078,741 bp.

#### Contig scaffolding

Though the five Columbia chromosomes were represented by only 10 contigs, we used reference-guide scaffolding to order and orient contigs, assign chromosome labels, and orient pseudomolecules to match the orientation of TAIR10 chromosomes. We ran RagTag (v1.0.1, --debug --aligner=nucmer --nucmer-params=’--maxmatch -l 100 -c 500’) using TAIR10 as the reference genome, but excluding ChrC and ChrM (-e) (*52, 53*). Three small contigs (3,200, 90,237 and 8,728 bp) consisting of low complexity sequence were not ordered and oriented and were removed from the assembly. After scaffolding, the 131,388,895 bp assembly was represented in five pseudomolecules corresponding to the five chromosomes of the Columbia genome. Chromosome 1 was gapless, while the other chromosomes contained one to four 100 bp gaps each (9 in total).

#### Pseudomolecule polishing and gap filling

We corrected mis-assemblies and filled gaps in the Columbia pseudomolecules with two rounds of Medaka (v1.2.1) ONT polishing (https://github.com/nanoporetech/medaka). For the first round of polishing, we aligned R9 ONT reads (>=50 kbp) to the pseudomolecules with mini_align (minimap2 v2.17-r941, -m). To avoid overcorrection in the centromere satellite sequences, we performed ‘marker-assisted filtering’ to remove alignments not anchored in putatively unique sequences (*5*) (https://github.com/malonge/T2T-Polish). We defined ‘marker’ *k*-mers as 21-mers that occurred once in the assembly and between 14 and 46 times (inclusive) in the Illumina reads. The first round of polishing was completed using ‘medaka consensus’ (--model r941_min_high_g360 --batch_size 200) and ‘medaka stitch’. The second round of polishing was performed as for the first round, except we aligned all R10 reads instead of R9 reads and the ‘medaka consensus’ model was set to “r103_min_high_g360”. As a result of ONT polishing, the assembly improved from a QV of 32.38 to 33.17 and 34.12 after the first and second rounds, respectively (*18*). After medaka polishing, the assembly contained only a single gap on chromosome 2.

Long-read ONT polishing was followed by short-read polishing of non-centromeres with DeepVariant (*54*). We first aligned Columbia genomic DNA Illumina reads to the pseudomolecules with bwa mem (v0.7.17-r1198-dirty) and we compressed and sorted alignments with samtools (v1.10) (*55, 56*). We then created a VCF file of potential polishing edits with DeepVariant (v1.1.0, -- model_type=WGS),“bcftools view” (v1.11, -e ’type=“ref”’ -i ’QUAL>1 && (GT=“AA” || GT=“Aa”)’ ) and “bcftools norm”. To avoid error-prone short-read polishing in the centromeres, we used Bedtools to remove polishing edits within the centromeres and we used BCFtools to derive a final consensus FASTA file (*57, 58*). Though short-read polishing did not alter the centromeres, it improved the overall assembly QV to 41.4616.

#### Telomere patching

We locally re-assembled and patched telomeric sequences for the 8 Columbia telomeres not adjacent to NORs (all except the beginning of chromosomes 2 and 4). We aligned all R9 reads to the TAIR10 reference with Winnowmap (v1.11, k=15, --MD -ax map-ont) and for each telomere, we collected all reads that aligned once to within 50 bp of the chromosome terminus (*9*). Using Bowtie (*59*) (v1.3.0, -S --all -v 0), we counted the occurrences of the telomeric repeat motif (‘CCCTAAA’) in each read, and the read with the most occurrences was designated as the ‘reference’ and all other reads were designated as the ‘query’. Local re-assembly was completed by aligning the query reads to the reference read and computing a consensus with ‘medaka_consensus’ (v1.2.1, -m r941_min_high_g360). To patch these telomere consensus sequences into the Columbia pseudomolecules, we identified the terminal BAC sequences for each of the 8 chromosome arms. For each chromosome arm, we aligned the terminal BAC sequence to the Columbia pseudomolecules and the telomere consensus sequence with Nucmer (v3.1, --maxmatch). Using these alignment coordinates, the consensus sequences were manually patched such that everything after the terminal BAC sequence was replaced with telomere consensus sequence. Telomeres were then manually confirmed to be structurally valid.

#### Assembly curation and preparation

After polishing and telomere patching, we performed final curation steps to correct lingering mis-assemblies and screen for contamination. First, while it was not straightforward to fill the remaining chromosome 2 gap *de novo*, we were able to replace the gap locus with the corresponding region in TAIR10. We found two BAC sequences flanking the gap locus that aligned concordantly to both the Col-0 pseudomolecules and TAIR10. These BAC contigs were aligned to the pseudomolecules and TAIR10 with Nucmer (v3.1, --maxmatch -l 250 -c 500) and the gap locus between the BAC contigs in the Columbia pseudomolecules was replaced with the corresponding TAIR10 locus between the BAC contigs.

To identify and correct structural mis-assemblies, we aligned Columbia ONT reads to the Columbia pseudomolecules and called structural variants (SVs). First, we used Bedtools ‘random’ (v2.29.2, -l 100000 -n 50000 -seed 23) to simulate 50,000 100 kbp exact reads from TAIR10. These reads, along with R9 (>=50 kbp) and R10 Columbia reads were aligned to the Columbia pseudomolecules with Winnowmap (v1.11, k=15, “--MD -ax map-pb” for TAIR10 reads and “--MD -ax map-ont” for ONT reads). After compressing and sorted alignments with samtools (v1.10), Sniffles (v1.0.12, -d 100 -n -1 -s 3) was used to infer SVs from each of the alignments (*60*). SVs with fewer than 30% of reads supporting the ALT allele were removed and the three resulting VCF files were merged with Jasmine (v1.0.10, max_dist=500 spec_reads=3 --output_genotypes) (*61*). There were a total of three variants called by all three read sets, including two deletions and one insertion that we corrected. REF and ALT alleles for these SVs were manually refined and validated, and ALT alleles were incorporated into the pseudomolecules using ‘bcftools consensus’.

Next, we manually inspected all gaps filled by Medaka and found that a 181 bp region containing a 100 bp gap on chromosome 5 was incorrectly replaced with 103 bp of sequence and we manually replaced the filled sequence with the original gap locus. Finally, we used VecScreen to do a final contamination screen. We first aligned the Columbia pseudomolecules to the VecScreen database with blastn (v2.5.0, -task blastn -reward 1 -penalty -5 -gapopen 3 -gapextend 3 -dust yes -soft_masking true -evalue 700 - searchsp 1750000000000 -outfmt “6 std score”). The BLAST alignments did not yield any ‘moderate’ or ‘strong’ matches to the database, so we determined that there was no contamination.

The final assembly contained five pseudomolecules with a single gap on chromosome 5, two missing telomeres, and partially resolved NOR sequences at the beginning of chromosomes 2 and 4. Chromosomes 1 and 3 were gapless and were completely sequence resolved from telomere-to-telomere (T2T). The final Col-CEN assembly FASTA file includes these 5 pseudomolecules and the Columbia chloroplast and mitochondria reference genomes.

#### Genome annotation

Genes were lifted-over from TAIR10 with Liftoff (v1.5.1, -copies -a 1 -s 1) (*62*). Since ChrC and ChrM were directly copied from TAIR10, their lift-over genes were replaced with their original TAIR10 annotations. We then used EDTA (v1.9.6, --sensitive 1 --anno 1 --evaluate 1) to perform *de novo* transposable element (TE) annotation, providing transcripts with “--cds” and the TAIR10 TE library with “--curatedlib” (*63, 64*). The TE annotation was supplemented with further manual annotation of the centromeric *ATHILA* elements (see section below). We used LASTZ to identify regions with similarity to *5S*, *45S* rDNA and the mitochondrial genome. To generate similarity heatmaps, the centromere region was divided into adjacent 5 kbp regions, which were compared using the pairwiseAlignment (type=’global’) and pid functions in R, using the Biostrings library. Sequences were compared in forward and reverse directions, and the highest percent sequence identity value kept. These values were then plotted in the heatmap.

#### *CEN180* repeat annotation

To identify repetitive regions, we divided the genome assembly into adjacent 1 kbp windows. In each window, for each position, we defined 12-mers and exactly matched these sequences to the rest of the window. We identified windows where the proportion of non-unique 12-mers was greater than 10% and merged contiguous windows that were above this threshold. For each region, we generated a histogram of the distances between 12-mers to test for periodic repeats. For example, if a region contains an arrayed tandem repeat of monomer size N, then a histogram of the 12-mer distances will show peaks at values N, N×2, N×3 … . The N value was obtained for each region, using the most frequent 12-mer distance. Next, 5 sequences of length N were randomly chosen from within the region and matched back to the sequence using the R function matchPattern (max.mismatch=N/3 with.indels=T). For each set of matches we identified overlapping repeats. If the overlap was less than 10 nucleotides, the overlap was divided at the midpoint between the repeats. If the overlap was 10 nucleotides or greater, the larger repeat was kept. The set of non-overlapping matches with the highest number was kept for further analysis. These sequence matches were aligned using mafft (--retree 2 --inputorder) (*65*), and a consensus repeat monomer was derived from the multiple sequence alignment. This consensus sequence was matched back to the region using matchPattern (max.mismatch=N/3 with.indels=T), and overlaps were treated in the same way.

Our approach identified 66,129 *CEN180* repeats with a mean length of 178 bp. The set of unique *CEN180* sequences (n=25,192) were aligned using mafft (--sparsescore 1000 --inputorder) (*65*). A consensus sequence was generated from the multiple sequence alignment, which was: 5′- AGTATAAGAACTTAAACCGCAACCCGATCTTAAAAGCCTAAGTAGTGTTTCCTTGTTAGAAGACACAAAGCCAAAGACTCATATGGACTTTGGCTACACCATGAAAGCTTTGAGAAGCAAGAAGAAGGTTGGTTAGTGTTTTGGAGTCGAATATGACTTGATGTCATGTGTATGATTG- 3′. In order to analyze *CEN180* diversity, for each position of the multiple sequence alignment (968 positions), we calculated the proportion of A, T, G, C and gaps. The alignment for each monomer at each position was then compared to these proportions and used to calculate a single nucleotide variant (SNV) score for the monomer. For example, if a monomer had an A in the alignment at a given position, and the overall proportion of A at that position was 0.7, the SNV score for that monomer would increase by 1-0.7. This was repeated for each position of the alignment, for each monomer. This ‘weighted’ SNV score was used to assess how similar a given *CEN180* monomer is to the genome-wide consensus. Alternatively, to compare pairwise differences between two specific monomers, the two sequences were compared along the length of the multiple sequence alignment and each instance of disagreement counted to give a ‘pairwise’ SNV score.

To identify higher order repeats (HORs) in a head-to-tail (tandem) orientation, each monomer was taken in turn and compared to all others using a matrix of pairwise SNV scores. If a pair of monomers had an SNV score of 5 or less, and were on the same strand, they were considered a match. For each match, monomers were extended by +1 unit in the same direction on the chromosome, and these were again compared for pairwise SNVs. This process was repeated until the next monomers had a pairwise SNV score higher than threshold, or the repeats were on opposite strands, or the end of the array was reached, with these conditions defining the end of the HOR. We also searched for repeats in head-to-head (inverted) orientation, which was identical apart from that repeats must be on opposite strands, and when monomers are extended to search for HORs, one is extended +1 position along the chromosome, whereas the other decreases -1. HORs were defined for each instance of 3 or more consecutive monomer matches. We define each HOR as consisting of block1 and block2 of *CEN180* monomers. The size of each block was recorded, in terms of monomers and base pairs, in addition to the distance between the block start coordinates. Cumulative pairwise SNVs per *CEN180* monomer were also calculated between each pair of blocks to provide a ‘block’ SNV score. To measure higher order repetition of each monomer, we summed the HOR block sizes in monomers, such that if a monomer was represented in three 5-mer blocks, it would score 15.

#### *ATHILA* annotation

To manually resolve the sequence of the centromeric *ATHILA* elements, we used LTRharvest (*66*) to complement the EDTA run that was used for the annotation of all Arabidopsis TEs (see above). We ran LTRharvest three times using ‘normal’, ‘strict’ and ‘very strict’ parameters. The parameters were gradually adjusted to allow us to capture the full-length sequence of the *ATHILA* family, based on older studies that reported the total and LTR lengths of intact *ATHILA* elements (*20*). These parameters were -maxlenltr 2500 -minltrlen 400 -mindistltr 2000 -maxdistltr 20000 -similar 75 -mintsd 0 -motif TGCA -motifmis 1 for the ‘normal’ run; -maxlenltr 2000 -minlenltr 1000 -mindistltr 4000 -maxdistltr 16000 - similar 80 -mintsd 3 -motif TGCA -motifmis 1 for the ‘strict’ run; and -maxlenltr 2100 -minlenltr 1100 -mindistltr 5000 -maxdistltr 14000 -similar 85 -mintsd 4 -motif TGCA -motifmis 1 -vic 20 for the ‘very strict’ run. Coordinates of predicted full-length elements from EDTA, LTRharvest and the manual dotplot annotation of centromeric TEs were merged and sequences aligned using MAFFT (*67*). Through these steps, we were able to define with base-pair resolution the external junctions of every *ATHILA* element, together with flanking sequences and the junctions of the LTRs with the internal domain (5′- LTR with PBS; PPT with 3′-LTR). Overall, we identified 46 intact elements (of which 34 have a detectable target site duplication), 5 fragmented *ATHILA* and 10 solo LTRs. We further identified open reading frames (minimum 150 nt) in the internal domain of the 46 intact elements using getorf in EMBOSS (*68*), and by running HMMER v3.3.2 (http://hmmer.org/) (-E 0.05 --domE 0.05) using a collection of Hidden Markov Models (HMMs) downloaded from Pfam (http://pfam.xfam.org/) that describe the coding domains of Gypsy retrotransposons: PF03732 for gag; PF13650, PF08284, PF13975 and PF09668 for protease; PF00078 for reverse transcriptase; PF17917, PF17919 and PF13456 for RNase-H; PF00665, PF13683, PF17921, PF02022, PF09337 and PF00552 for integrase. *ATHILA* elements may also contain an envelope-like ORF (*20*). To identify this domain (and because there is no HMM in Pfam that describes envelope-like genes of LTR retrotransposons), we used a previously published HMM developed by one of the co-authors (*69*).

#### ONT DNA methylation analysis

To identify methylation in CG, CHG and CHH sequence contexts we used DeepSignal-plant (v. 0.1) (*22*), which uses a deep-learning method based on bidirectional recurrent neural network (BRNN) with long short-term memory (LSTM) units to detect DNA 5mC methylation. R9 reads were filtered for length and accuracy using Filtlong (v0.2.0) (--min_mean_q 95, --min_length 30000. https://github.com/rrwick/Filtlong). Basecalled read sequence was annotated onto corresponding .fast5 files, and re-squiggled using Tombo (v 1.5.1). Methylation prediction for the CG, CHG, and CHH contexts were called using DeepSignal-plant using the respective models: model.dp2.CG.arabnrice2-1_R9.4plus_tem.bn13_sn16.balance.both_bilstm.b13_s16_epoch6.ckpt, model.dp2.CHG.arabnrice2- 1_R9.4plus_tem.bn13_sn16.denoise_sig1nal_bilstm.both_bilstm.b13_s16_epoch4.ckpt model.dp2.CHH.arabnrice2- 1_R9.4plus_tem.bn13_sn16.denoise_signal_bilstm.both_bilstm.b13_s16_epoch7.ckpt. The script call_modification_frequency.py provided in the DeepSignal-plant package was then used to generate the methylation frequency at each CG, CHG and CHH site. To identify CG context DNA methylation in Nanopore reads we also used Nanopolish (v 0.13.2), which uses a Hidden Markov model on the nanopore current signal to distinguish 5-methylcytosine from unmethylated cytosine. Reads were first filtered for length and accuracy using Filtlong (v0.2.0) (--min_mean_q 95, --min_length 15000. https://github.com/rrwick/Filtlong). The subset was then indexed to the fast5 files and aligned to the genome using Winnowmap (v1.11, -ax map-ont). The read fastq, alignment bam, and fast5 files were used as an input to the Nanopolish call-methylation function. The script calculate_methylation_frequency.py provided in the Nanopolish package was then used to generate the methylation frequency at each CG containing *k*-mer.

#### ChIP-seq and MNase-seq data alignment and processing

Deduplicated paired-end ChIP-seq and MNase-seq reads **(Table S5)** were processed with Cutadapt v1.18 to remove adapter sequences and low-quality bases (Phred+33-scaled quality <20) (*70*). Trimmed reads were aligned to the Col-CEN genome assembly using Bowtie2 v2.3.4.3 with the following settings: --very-sensitive --no-mixed --no-discordant -k 10 (*71*). Up to 10 valid alignments were reported for each read pair. Read pairs with Bowtie2-assigned MAPQ <10, including those that aligned equally well to more than one location, were discarded using Samtools v1.9 (*56*). For retained read pairs that aligned to multiple locations, with varying alignment scores, the best alignment was selected. Alignments with more than 2 mismatches or consisting of only one read in a pair were discarded. Single-end SPO11-1-oligo reads were processed and aligned to the Col-CEN assembly using an equivalent pipeline without paired-end options, as described (*30*). For each data set, bins per million mapped reads (BPM; equivalent to transcripts per million, TPM, for RNA-seq data) coverage values were generated in bigWig and bedGraph formats with the bamCoverage tool from deepTools v3.1.3 (*72*). Reads that aligned to chloroplast or mitochondrial DNA were excluded from this coverage normalization procedure.

#### RNA-seq data alignment and processing

Paired-end RNA-seq reads (2×100 bp) were processed with Trimmomatic v0.38 to remove adapter sequences and low-quality bases (Phred+33-scaled quality <3 at the beginning and end of each read, and average quality <15 in 4-base sliding windows) **(Table S5)** (*30, 73*). Trimmed reads were aligned to the Col-CEN genome assembly using STAR v2.7.0d with the following settings: -- outFilterMultimapNmax 100 --winAnchorMultimapNmax 100 --outMultimapperOrder Random -- outFilterMismatchNmax 2 --outSAMattributes All --twopassMode Basic --twopass1readsN -1 (*74*). Read pairs with STAR-assigned MAPQ <3 were discarded using Samtools v1.9 (*56*). For retained read pairs that aligned to multiple locations, with varying alignment scores, the best alignment was selected. Alignments with more than 2 mismatches, or consisting of only one read in a pair, were discarded.

#### Small RNA-seq data alignment and processing

Small RNA-seq reads **(Table S5)** (*31*), were processed with BBDuk from BBMap v38.22 (*75*), to remove ribosomal sequences and Cutadapt v1.18 (*70*) to remove adapter sequences and low-quality bases (Phred+33-scaled quality <20). Trimmed reads were aligned to the Col-CEN genome assembly using Bowtie v1.2.2, allowing no mismatches (*59*). For reads that aligned to multiple locations, with varying alignment scores, the best alignment was selected. For each small RNA size class (18–26 nucleotides), TPM values in adjacent genomic windows were calculated based on the total retained alignments (across all size classes) in the library.

#### Bisulfite sequencing data alignment and processing

Paired-end bisulfite sequencing reads (2×90 bp) **(Table S5)** (*31, 76*), were processed with Trim Galore v0.6.4 to remove sequencing adapters, low-quality bases (Phred+33-scaled quality <20) and 3 bases from the 5′ end of each read (*77*). Trimmed reads were aligned to the Col-CEN assembly Bismark v0.20.0 (*78*). Read pairs that aligned equally well to more than one location and duplicate alignments were discarded. Methylated cytosine calls in CG, CHG and CHH sequence contexts were extracted and context-specific DNA methylation proportions were generated in bedGraph and bigWig formats using the bismark2 bedGraph and UCSC bedGraphToBigWig tools.

#### Fine-scale profiling around feature sets

Fine-scale profiles around *CEN180* (*n*=66,129), randomly positioned loci of the same number and width distribution (*n*=66,129), centromeric *ATHILA* elements (*n*=46), and non-centromeric *GYPSY* elements (*n*=3,980) were calculated for ChIP-seq, MNase-seq, RNA-seq, small RNA-seq and bisulfite-seq data sets by providing the above-described bigWig files to the computeMatrix tool from deepTools v3.1.3 in ‘scale-regions’ mode (*72*). Each feature was divided into non-overlapping, proportionally scaled windows between start and end coordinates, and flanking regions were divided into 10 bp windows. Mean values for each data set were calculated within each window, generating a matrix of profiles in which each row represents a feature with flanking regions and each column a window. Coverage profiles for a ChIP input sequencing library and a gDNA library **(Table S5)** were used in conjunction with those for ChIP-seq and SPO11-1-oligo libraries, respectively, to calculate windowed log_2_([ChIP+1]/[control+1]) coverage ratios for each feature. Meta-profiles (windowed means and 95% confidence intervals) for each group of features were calculated and plotted using the feature profiles in R version 4.0.0.

#### Crossover mapping

Total data from 96 Col×Ler genomic DNA F_2_ sequencing libraries (2×150 bp) were aligned to the Col-CEN assembly using bowtie2 (default settings), which gave 87.15% overall alignment. Polymorphisms were identified using the alignment files with samtools mpileup (-vu -f) and bcftools call (-mv -Oz). The resulting polymorphisms were filtered for SNPs (n=522,931), which was used as the ‘complete’ polymorphism set in TIGER (*79*). These SNPs were additionally filtered by, (i) removing SNPs with a quality score less than 200, (ii) removing SNPs where total coverage was greater than 300, or less than 50 (mean coverage=170.8), (iii) removing SNPs that had reference allele coverage less than 20 or greater than 150, (iv) removing SNPs that had variant allele coverage greater than 130, (v) masking SNPs that overlapped transposon and repeat annotations and (vi) masking SNPs within the main *CEN180* arrays. This resulted in a ‘filtered’ set of 171,947 SNPs for use in TIGER. DNA sequencing data from 260 wild type Col×Ler F_2_ genomic DNA (192 from ArrayExpress E-MTAB-4657 and 68 from E-MTAB-6577) was aligned to the Col-CEN assembly using bowtie2 (default settings) and the alignment analyzed at the previously defined ‘complete’ SNPs using samtools mpileup (-vu -f) and bcftools call (-m -T). These sites were used as an input to TIGER, which identifies crossover positions by genotype transitions (*79*). A total of 2,042 crossovers were identified with a mean resolution of 1,011 bp.

#### Epitope tagging of *V5-DMC1*

The *DMC1* promoter region was PCR amplified from Col-0 genomic DNA using the Dmc1-PstI-fw and Dmc1-SphI-rev oligonucleotides **(Table S6)**. The remainder of the *DMC1* promoter, gene and terminator were amplified with oligonucleotides Dmc1-SphI-fw and Dmc1-NotI-rev. The resulting PCR fragments were digested with *Pst*I and *Sph*I, or *Sph*I and *Not*I, respectively, and cloned into *Pst*I- *Not*I-digested pGreen0029 vector to yield a pGreen-DMC1 construct. To insert 3 N-terminal V5 epitope tags, first two fragments were amplified with DMC1-Nco-F and 3N-V5-R and 3N-V5-F and Dmc1-Spe-rev and then used in an overlap PCR reaction using the DMC1-Nco-F and Dmc1-Spe-rev oligonucleotides. The PCR product resulting from the overlap PCR was digested with *Nco*I and *Spe*I and cloned into *Nco*I- and *Spe*I-digested pGreen-DMC1. The resulting binary vector was used to transform *dmc1-3/+* heterozygotes (SAIL_126_F07). We used dmc1-seq11 and Dmc1-Spe-rev oligonucleotides to amplify wild type *DMC1* allele and Dmc1-Spe-rev and LA27 to amplify the *dmc1-3* T-DNA mutant allele. The presence of the *V5-DMC1* transgene was detected with N-screen-F and N-screen-R oligonucleotides. This oligonucleotide pair amplifies a 74 bp product in Col and a 203 bp product in *V5-DMC1*. To identify *dmc1-3* homozygotes in the presence of *V5-DMC1* transgenes, we used DMC1-genot-compl-F and DMC1-genot-compl-R oligonucleotides, which allowed us to distinguish between the wild type *DMC1* gene and *V5-DMC1* transgene. All oligonucleotide sequences are provided in **Table S6**.

#### Immunocytological analysis

Fresh buds at floral stage 8 and 9 were dissected to release the anthers that contain male meiocytes (*80*). Chromosome spreads of meiotic and mitotic cells from anthers were performed, followed by immunofluorescent staining of proteins, as described (*28*). The antibodies used in this study were: α-ZYP1 (rabbit, 1/500 dilution) (*81*), α-H3K9me2 (mouse, 1/200 dilution) (Abcam, ab1220), α-CENH3 (rabbit, 1/100 dilution) (Abcam, ab72001) and α-V5 (chicken, 1/200 dilution) (Abcam, ab9113). Chromosomes stained with ZYP1, CENH3 and H3K9me2 were visualized with a DeltaVision Personal DV microscope (Applied Precision/GE Healthcare). Chromosomes stained with DMC1-V5 and CENH3 were visualized with a Leica SP8 confocal microscope. Chromosomes stained with H3K9me2 were visualized with a Stimulated emission depletion nanoscopy mounted on an inverted IX71 Olympus microscope.

---

**Table S1.**
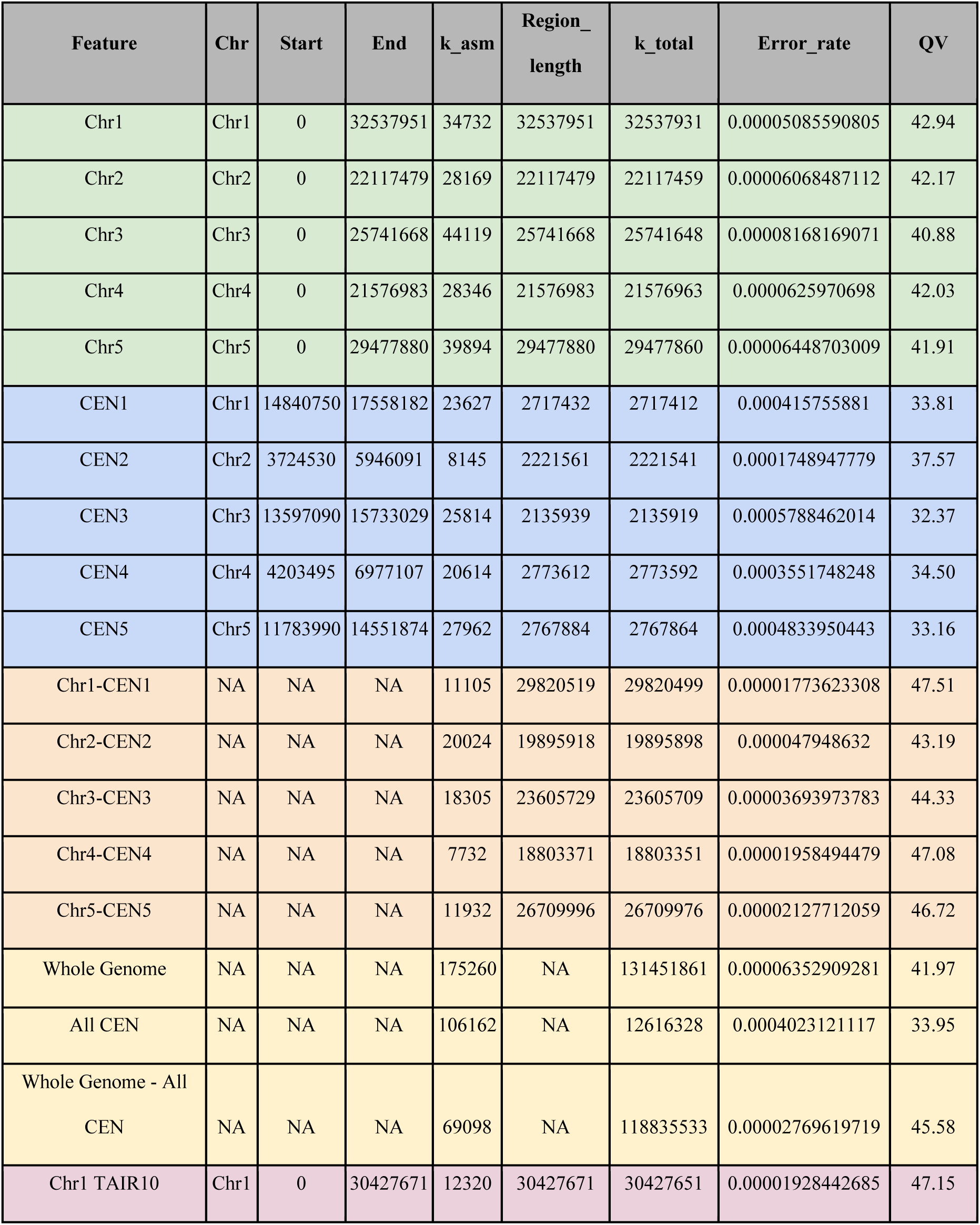
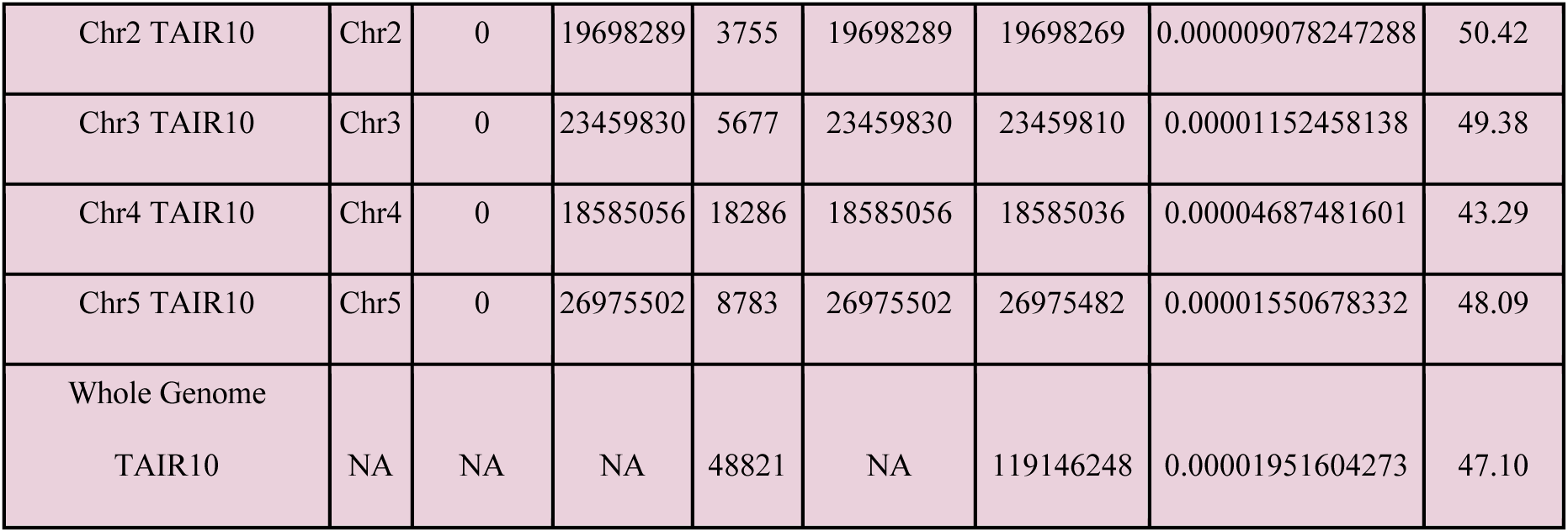
Consensus quality (QV) score of Arabidopsis genome assemblies. Consensus quality scores (QV) were calculated from ‘missing’ 21-mers (k_asm) present in the Col-CEN assembly, but not present in the short read Illumina library. k_total shows the total number of 21-mers. QV scores were calculated for Col-CEN individual chromosomes (green), centromeres (blue), chromosome arms (orange), or the whole genome (yellow). QV scores for TAIR10 are shown for comparison (pink).

| Feature | Chr | Start | End | k_asm | Region_<br>length | k_total | Error_rate | QV |
| --- | --- | --- | --- | --- | --- | --- | --- | --- |
| Chr1 | Chr1 | 0 | 32537951 | 34732 | 32537951 | 32537931 | 0.00005085590805 | 42.94 |
| Chr2 | Chr2 | 0 | 22117479 | 28169 | 22117479 | 22117459 | 0.00006068487112 | 42.17 |
| Chr3 | Chr3 | 0 | 25741668 | 44119 | 25741668 | 25741648 | 0.00008168169071 | 40.88 |
| Chr4 | Chr4 | 0 | 21576983 | 28346 | 21576983 | 21576963 | 0.0000625970698 | 42.03 |
| Chr5 | Chr5 | 0 | 29477880 | 39894 | 29477880 | 29477860 | 0.00006448703009 | 41.91 |
| CEN1 | Chr1 | 14840750 | 17558182 | 23627 | 2717432 | 2717412 | 0.000415755881 | 33.81 |
| CEN2 | Chr2 | 3724530 | 5946091 | 8145 | 2221561 | 2221541 | 0.0001748947779 | 37.57 |
| CEN3 | Chr3 | 13597090 | 15733029 | 25814 | 2135939 | 2135919 | 0.0005788462014 | 32.37 |
| CEN4 | Chr4 | 4203495 | 6977107 | 20614 | 2773612 | 2773592 | 0.0003551748248 | 34.50 |
| CEN5 | Chr5 | 11783990 | 14551874 | 27962 | 2767884 | 2767864 | 0.0004833950443 | 33.16 |
| Chr1-CEN1 | NA | NA | NA | 11105 | 29820519 | 29820499 | 0.00001773623308 | 47.51 |
| Chr2-CEN2 | NA | NA | NA | 20024 | 19895918 | 19895898 | 0.000047948632 | 43.19 |
| Chr3-CEN3 | NA | NA | NA | 18305 | 23605729 | 23605709 | 0.00003693973783 | 44.33 |
| Chr4-CEN4 | NA | NA | NA | 7732 | 18803371 | 18803351 | 0.00001958494479 | 47.08 |
| Chr5-CEN5 | NA | NA | NA | 11932 | 26709996 | 26709976 | 0.00002127712059 | 46.72 |
| Whole Genome | NA | NA | NA | 175260 | NA | 131451861 | 0.00006352909281 | 41.97 |
| All CEN | NA | NA | NA | 106162 | NA | 12616328 | 0.0004023121117 | 33.95 |
| Whole Genome - All<br>CEN | NA | NA | NA | 69098 | NA | 118835533 | 0.00002769619719 | 45.58 |
| Chr1 TAIR10 | Chr1 | 0 | 30427671 | 12320 | 30427671 | 30427651 | 0.00001928442685 | 47.15 |
| Chr2 TAIR10 | Chr2 | 0 | 19698289 | 3755 | 19698289 | 19698269 | 0.000009078247288 | 50.42 |
| Chr3 TAIR10 | Chr3 | 0 | 23459830 | 5677 | 23459830 | 23459810 | 0.00001152458138 | 49.38 |
| Chr4 TAIR10 | Chr4 | 0 | 18585056 | 18286 | 18585056 | 18585036 | 0.00004687481601 | 43.29 |
| Chr5 TAIR10 | Chr5 | 0 | 26975502 | 8783 | 26975502 | 26975482 | 0.00001550678332 | 48.09 |
| Whole Genome<br>TAIR10 | NA | NA | NA | 48821 | NA | 119146248 | 0.00001951604273 | 47.10 |

**Table S2.** Unique and repeated CEN180 monomer sequences within and between chromosomes. CEN180 monomers were compared across the genome to identify unique versus repeated sequences. For repeated sequences we show which chromosomes they occurred on.

| Chr | Total | Unique | Repeated | Chr1 | Chr2 | Chr3 | Chr4 | Chr5 | Chr2,<br>Chr4, Chr5 |
| --- | --- | --- | --- | --- | --- | --- | --- | --- | --- |
| Chr1 | 13,802 | 4,978 | Chr1 | 8,572 | 0 | 250 | 0 | 2 | 25 |
| Chr2 | 12,297 | 4,271 | Chr2 |  | 7,979 | 20 | 20 | 7 |  |
| Chr3 | 11,623 | 4,637 | Chr3 |  |  | 6,979 | 0 | 7 |  |
| Chr4 | 15,610 | 5,418 | Chr4 |  |  |  | 10,192 | 0 |  |
| Chr5 | 12,797 | 5,856 | Chr5 |  |  |  |  | 6,916 |  |
| All | 66,129 | 25,160 | Total | 8,572 | 7,979 | 7,249 | 10,212 | 6,932 | 40,969 |

**Table S3.** *CEN180* higher order repeats. *CEN180* monomers were classified as being the same if they shared 5 or fewer SNVs, and consecutive blocks identified as higher order repeats (HORs). HORs are in a tandem orientation and are classified as being intra- or inter-chromosome. The mean HOR block size, in monomers and bp, and the mean distance between intra-chromosome HORs (bp) are listed.

| Chr | Monomers | Intra-chromosome HORs | Inter-chromosome HORs | Mean HOR monomers | Mean HOR block (bp) | Mean HOR distance (bp) |
| --- | --- | --- | --- | --- | --- | --- |
| 1 | 13,802 | 154,846 | 40 | 3.63 | 646 | 210,272 |
| 2 | 12,297 | 124,095 | 86 | 3.55 | 631 | 308,877 |
| 3 | 11,623 | 84,531 | 1 | 3.89 | 693 | 166,749 |
| 4 | 15,610 | 118,671 | 0 | 3.64 | 647 | 329,844 |
| 5 | 12,797 | 18,690 | 0 | 4.41 | 783 | 77,533 |
| All | 66,129 | 500,833 | 127 | 3.69 | 656 | 250,737 |

**Table S4.** Centromeric ATHILA retrotransposons. Analysis of 69 gaps in the *CEN180* arrays identified 46 intact and five fragmented *ATHILA* retrotransposons, as well as 10 solo LTRs. For each sequence we report the subfamily class based on the TAIR10 classification, and information on length, strand, target site duplications (TSDs), long terminal repeat (LTR) positions, LTR % identity, the the 5′-LTR/primer binding site (PBS) and 3′-polypurine tract (PPT)/LTR junctions, and hits with Hidden Markov Models (HMMs) that describe Gypsy retrotransposon open reading frames (a six digit identifier is provided, where 1 indicates the presence of (i) gag, (ii) protease, (iii) reverse transcriptase, (iv) RNaseH, (v) integrase or (vi) envelope genes). The structure column indicates whether a transposon was full-length (i.e. it contained clearly identified LTRs and PBS/PPT junctions with the internal domain), fragmented (i.e. when one or both LTRs were missing or when a complex nested insertion was found), or a solo LTR. In addition to the *ATHILA* elements, a small number of other TEs were identified, but not further analyzed due to their fragmented or nested organization. These elements are shown at the end of the Table. Note that for these elements the coordinates refer to the position of the gaps and not the TEs within the gaps.

| Chr | Class | Start | End | Length (bp) | Strand | Left TSD | Right TSD | 5'-LTR | 3'-LTR | LTR % identity | 5'-LTR/PBS junction | PPT/3'-LTR junction | Structure | HMMs gag-pr-rt-rh-int-env |
| --- | --- | --- | --- | --- | --- | --- | --- | --- | --- | --- | --- | --- | --- | --- |
| 1 | *ATHILA2* | 14963269 | 14973475 | 10206 | - | AAGTT | AAGTT | 1695 | 1757 | 99.23 | 1694 | 8450 | full-length | 110011 |
| 2 | *ATHILA6A* | 4949814 | 4961120 | 11306 | + | TTACG | TTACG | 1804 | 1799 | 99.05 | 1803 | 9508 | full-length | 110011 |
| 2 | *ATHILA5* | 5188085 | 5198642 | 10557 | + | no TSD | no TSD | 1540 | 1539 | 99.74 | 1539 | 9019 | full-length | 110001 |
| 2 | *ATHILA6A* | 5440344 | 5451684 | 11340 | - | TTAAA | TTAAA | 1814 | 1810 | 99.67 | 1813 | 9531 | full-length | 110011 |
| 3 | *ATHILA2* | 13790673 | 13801819 | 11146 | + | GTACC | ATACC | 1805 | 1804 | 98.17 | 1804 | 9343 | full-length | 110010 |
| 3 | *ATHILA* | 14419926 | 14429549 | 9623 | - | no TSD | no TSD | 1293 | 1595 | 98.32 | 1292 | 8029 | full-length | 110001 |
| 3 | *ATHILA* | 14745694 | 14755319 | 9625 | - | no TSD | no TSD | 1293 | 1596 | 98.08 | 1292 | 8030 | full-length | 110001 |
| 4 | *ATHILA* | 5706402 | 5717133 | 10731 | - | CAAAG | CAAAG | 1654 | 1659 | 99.64 | 1653 | 9073 | full-length | 110010 |
| 4 | *ATHILA* | 5739753 | 5749900 | 10147 | + | TACTC | TACTC | 1205 | 1588 | 98.75 | 1204 | 8560 | full-length | 110010 |
| 4 | *ATHILA6B* | 6015319 | 6026902 | 11583 | + | no TSD | no TSD | 1807 | 1806 | 99.17 | 1806 | 9778 | full-length | 111111 |
| 4 | *ATHILA* | 6081947 | 6092675 | 10728 | + | no TSD | no TSD | 1655 | 1655 | 99.15 | 1654 | 9074 | full-length | 110010 |
| 4 | *ATHILA* | 6620303 | 6630820 | 10517 | + | no TSD | no TSD | 1548 | 1546 | 99.48 | 1547 | 8972 | full-length | 110010 |
| 4 | *ATHILA* | 6642167 | 6652842 | 10675 | + | no TSD | no TSD | 1652 | 1652 | 99.52 | 1651 | 9024 | full-length | 110010 |
| 4 | *ATHILA2* | 6673701 | 6684761 | 11060 | + | ATAAG | ATAAG | 1755 | 1761 | 98.92 | 1754 | 9300 | full-length | 110011 |
| 4 | *ATHILA2* | 6806885 | 6811170 | 4285 | - | no TSD | no TSD | 1168 | 1170 | 97.94 | 1167 | 3116 | full-length | 100010 |
| 4 | *ATHILA* | 6927958 | 6938157 | 10199 | - | ATATG | ATATG | 1553 | 1553 | 99.1 | 1552 | 8647 | full-length | 110000 |
| 5 | *ATHILA6A* | 11814343 | 11825681 | 11338 | - | CTTTT | CTTTT | 1803 | 1806 | 99.5 | 1802 | 9533 | full-length | 110111 |
| 5 | ATHILA6B | 11841351 | 11852931 | 11580 | - | AGACG | AGACG | 1806 | 1804 | 99 | 1805 | 9777 | full-length | 111111 |
| 5 | ATHILA6A | 11898313 | 11909640 | 11327 | + | CATAC | CATAC | 1804 | 1801 | 98.94 | 1803 | 9527 | full-length | 110111 |
| 5 | ATHILA2 | 11911707 | 11919267 | 7560 | + | no TSD | no TSD | 1517 | 1443 | 96.76 | 1516 | 6118 | full-length | 111100 |
| 5 | ATHILA2 | 11922096 | 11933394 | 11298 | - | GTATG | GTATG | 1726 | 1866 | 97.5 | 1725 | 9433 | full-length | 111011 |
| 5 | ATHILA6A | 11991440 | 12001550 | 10110 | + | no TSD | no TSD | 1806 | 567 | 99.47 | 1805 | 9544 | full-length | 110111 |
| 5 | ATHILA6B | 12090681 | 12102258 | 11577 | - | GGAGT | GTAGG | 1802 | 1806 | 99.11 | 1801 | 9772 | full-length | 111111 |
| 5 | ATHILA6B | 12309482 | 12321044 | 11562 | + | TATCC | TATCC | 1761 | 1804 | 99.37 | 1760 | 9759 | full-length | 111111 |
| 5 | ATHILA5 | 12459028 | 12469585 | 10557 | + | GTTGG | GTTGG | 1514 | 1539 | 99.27 | 1513 | 9019 | full-length | 110011 |
| 5 | ATHILA6A | 12482175 | 12493514 | 11339 | - | TTTTT | TTTTT | 1808 | 1801 | 98.28 | 1807 | 9539 | full-length | 110111 |
| 5 | ATHILA6A | 12559979 | 12571324 | 11345 | + | AGACA | AGACA | 1804 | 1796 | 99 | 1803 | 9550 | full-length | 110111 |
| 5 | ATHILA6A | 12662889 | 12674221 | 11332 | + | no TSD | no TSD | 1807 | 1801 | 99.39 | 1806 | 9532 | full-length | 110111 |
| 5 | ATHILA5 | 13217369 | 13227423 | 10054 | + | TTTGG | TTTGG | 1506 | 1535 | 99.2 | 1505 | 8520 | full-length | 110001 |
| 5 | ATHILA6A | 13343808 | 13355133 | 11325 | + | GAAGA | GAAGA | 1805 | 1800 | 99.17 | 1804 | 9526 | full-length | 110111 |
| 5 | ATHILA6A | 13743447 | 13754398 | 10951 | - | no TSD | no TSD | 1809 | 1470 | 99.05 | 1808 | 9482 | full-length | 110111 |
| 5 | ATHILA2 | 13832336 | 13843514 | 11178 | - | CTTTC | CTTTC | 1760 | 1557 | 97.81 | 1759 | 9622 | full-length | 110011 |
| 5 | ATHILA2 | 13851467 | 13864693 | 13226 | + | AAGCC | AAGCC | 1822 | 1771 | 96.71 | 1821 | 11456 | full-length | 111111 |
| 5 | ATHILA6B | 13864859 | 13876349 | 11490 | - | CAAGA | CAAGA | 1809 | 1802 | 98.94 | 1808 | 9689 | full-length | 111110 |
| 5 | ATHILA6A | 13893684 | 13904957 | 11273 | - | AGTTG | AGTTG | 1806 | 1805 | 99.28 | 1805 | 9469 | full-length | 110111 |
| 5 | ATHILA | 14034803 | 14045501 | 10698 | + | AAGAC | AAGAC | 1654 | 1652 | 99.15 | 1653 | 9047 | full-length | 110010 |
| 5 | ATHILA6B | 14074285 | 14085846 | 11561 | - | TTAGG | TTAGG | 1806 | 1804 | 98.67 | 1805 | 9758 | full-length | 111111 |
| 5 | ATHILA6A | 14142441 | 14153832 | 11391 | - | AACAC | AACAC | 1805 | 1803 | 98.5 | 1804 | 9589 | full-length | 110111 |
| 5 | ATHILA | 14222801 | 14234805 | 12004 | - | TTTTT | TCTTT | 1262 | 1613 | 95.11 | 1261 | 10392 | full-length | 110100 |
| 5 | ATHILA2 | 14253091 | 14264004 | 10913 | + | CGAGT | AGAGT | 1749 | 1749 | 98.11 | 1748 | 9165 | full-length | 110011 |
| 5 | ATHILA2 | 14278114 | 14289497 | 11383 | - | GTAGC | GTAGC | 1868 | 1713 | 90.27 | 1867 | 9671 | full-length | 110011 |
| 5 | ATHILA6A | 14291467 | 14302787 | 11320 | - | TCATT | TCATT | 1801 | 1791 | 99.05 | 1800 | 9530 | full-length | 110011 |
| 5 | ATHILA2 | 14387820 | 14399352 | 11532 | - | GACTC | GACTC | 1736 | 1749 | 95.63 | 1735 | 9784 | full-length | 101110 |
| 5 | ATHILA6B | 14400831 | 14412235 | 11404 | + | ACCAC | ACCGC | 1755 | 1766 | 94.61 | 1754 | 9639 | full-length | 111110 |
| 5 | ATHILA6A | 14420927 | 14431630 | 10703 | + | TGTAT | TGTAT | 1805 | 1167 | 99.23 | 1804 | 9537 | full-length | 110111 |
| 5 | ATHILA2 | 14515541 | 14529552 | 14011 | - | TAGCC | TAGCC | 1744 | 1746 | 97.99 | 1743 | 12266 | full-length | 111110 |
| 4 | ATHILA | 6564400 | 6569193 | 4793 | - | na | na | na | na | na | na | na | fragment | na |
| 4 | ATHILA2 | 6657901 | 6660658 | 2757 | + | na | na | na | na | na | na | na | fragment | na |
| 4 | ATHILA2 | 6666875 | 6669635 | 2760 | + | na | na | na | na | na | na | na | fragment | na |
| 5 | ATHILA6A | 14311956 | 14332997 | 21041 | + | CCATC | CCATC | na | na | na | na | na | fragment/<br>mosaic | na |
| 5 | ATHILA6B | 14461746 | 14482520 | 20774 | + | no TSD | no TSD | na | na | na | na | na | fragment | na |
| 1 | ATHILA2 | 14984261 | 14986017 | 1756 | - | AGT--T | AGTACT | na | na | na | na | na | solo LTR | na |
| 1 | ATHILA2 | 15006981 | 15008741 | 1760 | - | AGTACT | AGTACT | na | na | na | na | na | solo LTR | na |
| 4 | ATHILA | 5735848 | 5737435 | 1587 | + | TACTC | TACTC | na | na | na | na | na | solo LTR | na |
| 4 | ATHILA | 5764410 | 5766062 | 1652 | - | ACTCA | ACTCA | na | na | na | na | na | solo LTR | na |
| 5 | ATHILA6 | 12273020 | 12274825 | 1805 | - | TCTTC | TCTTC | na | na | na | na | na | solo LTR | na |
| 5 | ATHILA6 | 12521880 | 12523675 | 1795 | + | no TSD | no TSD | na | na | na | na | na | solo LTR | na |
| 5 | ATHILA5 | 12617002 | 12618543 | 1541 | + | no TSD | no TSD | na | na | na | na | na | solo LTR | na |
| 5 | *ATHILA5* | 12739533 | 12741074 | 1541 | + | no TSD | no TSD | na | na | na | na | na | solo LTR | na |
| 5 | *ATHILA4* | 14459130 | 14460352 | 1222 | - | ACATT | ACATT | na | na | na | na | na | solo LTR | na |
| 5 | *ATHILA6* | 14534797 | 14536744 | 1947 | - | GTCCG | GTTCG | na | na | na | na | na | solo LTR | na |

Non-*ATHILA* insertions:
| Chr | Class | Start | End | Length (bp) | Strand |
| --- | --- | --- | --- | --- | --- |
| 1 | *En/Spm* | 14975384 | 14983736 | 8352 | na |
| 1 | *AT3TE35825* | 15518346 | 15519532 | 1186 | + |
| 3 | *Gypsy* | 13704904 | 13711848 | 6944 | na |
| 4 | *ATGP2_AT2TE00075* | 4522011 | 4529406 | 7395 | - |
| 4 | *Gypsy – En/Spm* | 6903181 | 6912459 | 9278 | na |
| 5 | *DNA TE* | 12618893 | 12627936 | 9043 | na |
| 5 | *DNA TE* | 12741424 | 12750457 | 9033 | na |
| 5 | *unknown* | 14091855 | 14094574 | 2719 | + |

**Table S5.** Summary of short-read Illumina sequencing libraries aligned to the Col-CEN assembly.

| Library | Study accession | Run accession | Read length | Tissue | References |
| --- | --- | --- | --- | --- | --- |
| CENH3<br>ChIP-seq | PRJNA349052 | SRR4430537 | 2×100 bp | Seedling | (13) |
| H3K9me2<br>ChIP-seq | PRJEB36221 | ERR3813867 | 2×75 bp | Floral<br>bud | (28) |
| H3K27me1 ChIP-seq | PRJEB36221 | ERR3813864 | 2×75 bp | Floral<br>bud | (28) |
| H3K4me1<br>ChIP-seq | PRJEB36221 | ERR3813865 | 2×75 bp | Floral<br>bud | (28) |
| H3K4me2<br>ChIP-seq | PRJEB36221 | ERR3813866 | 2×75 bp | Floral<br>bud | (28) |
| H3K4me3<br>ChIP-seq | PRJEB15183 | ERR1590146 | 2×150 bp | Floral<br>bud | (30) |
| H3K27me3 ChIP-seq | PRJNA252965 | SRR1509478 | 2×100 bp | Floral<br>bud | (82) |
| H2A.W6<br>ChIP-seq | PRJNA377526 | SRR5298545 | 2×50 bp | Leaf | (83) |
| H2A.W7<br>ChIP-seq | PRJNA377526 | SRR5298546 | 2×50 bp | Leaf | (83) |
| H2A.Z ChIP-seq | PRJNA219442 | SRR988546 | 50 bp | Leaf | (25) |
| REC8<br>ChIP-seq | PRJEB36221 | ERR3813871 | 2×75 bp | Floral<br>bud | (28) |
| ASY1<br>ChIP-seq | PRJEB36320 | ERR3829803 | 2×75 bp | Floral<br>bud | <u>(29)</u> |
| SPO11-1-oligos | PRJEB15185 | ERR1590157 | 50 bp | Floral<br>bud | <u>(30)</u> |
| MNase-seq | PRJEB15184 | ERR1590154 | 2×100 bp | Floral<br>bud | <u>(30)</u> |
| ChIP input | PRJEB36221 | ERR3813873 | 2×75 bp | Floral<br>bud | <u>(28)</u> |
| ChIP input | PRJNA219442 | SRR988541 | 50 bp | Leaf | <u>(25)</u> |
| gDNA | PRJEB23842 | ERR2215865 | 2×100 bp | Floral<br>bud | <u>(30)</u> |
| RNA-seq (Col-0 and<br>*met1-3*) | PRJEB19033 | ERR1797989–<br>ERR1797990 | 2×100 bp | Floral<br>bud | <u>(30)</u> |
| Bisulfite-seq (Col-0) | PRJNA289907 | SRR2102617–<br>SRR2102619 | 2×90 bp | Seedling | <u>(76)</u> |
| Bisulfite-seq (Col-0 and<br>*met1-3*) | PRJEB9919 | ERR965674–<br>ERR965677 | 2×90 bp | Leaf | <u>(31)</u> |
| Small RNA-seq (Col-0<br>and *met1-3*) | PRJEB9919 | ERR966148–<br>ERR966149 | 50 bp | Leaf | <u>(31)</u> |
| Col×Ler genomic DNA<br>F<sub>2</sub> | E-MTAB-4657 E-MTAB-6577 | E-MTAB-4657 E-MTAB-6577 | 2×150 bp | Leaf | <u>(84, 85)</u> |

**Table S6.**
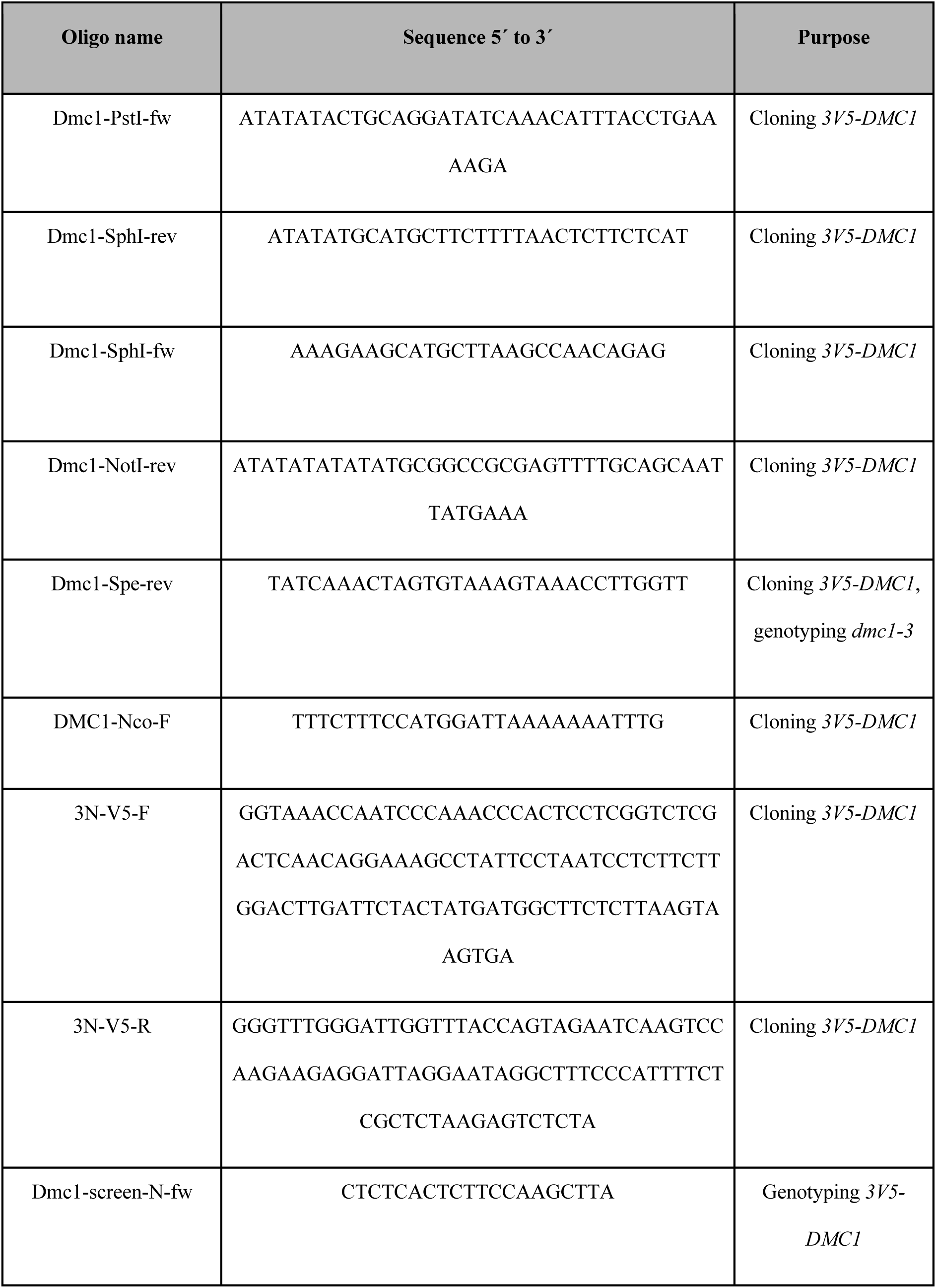
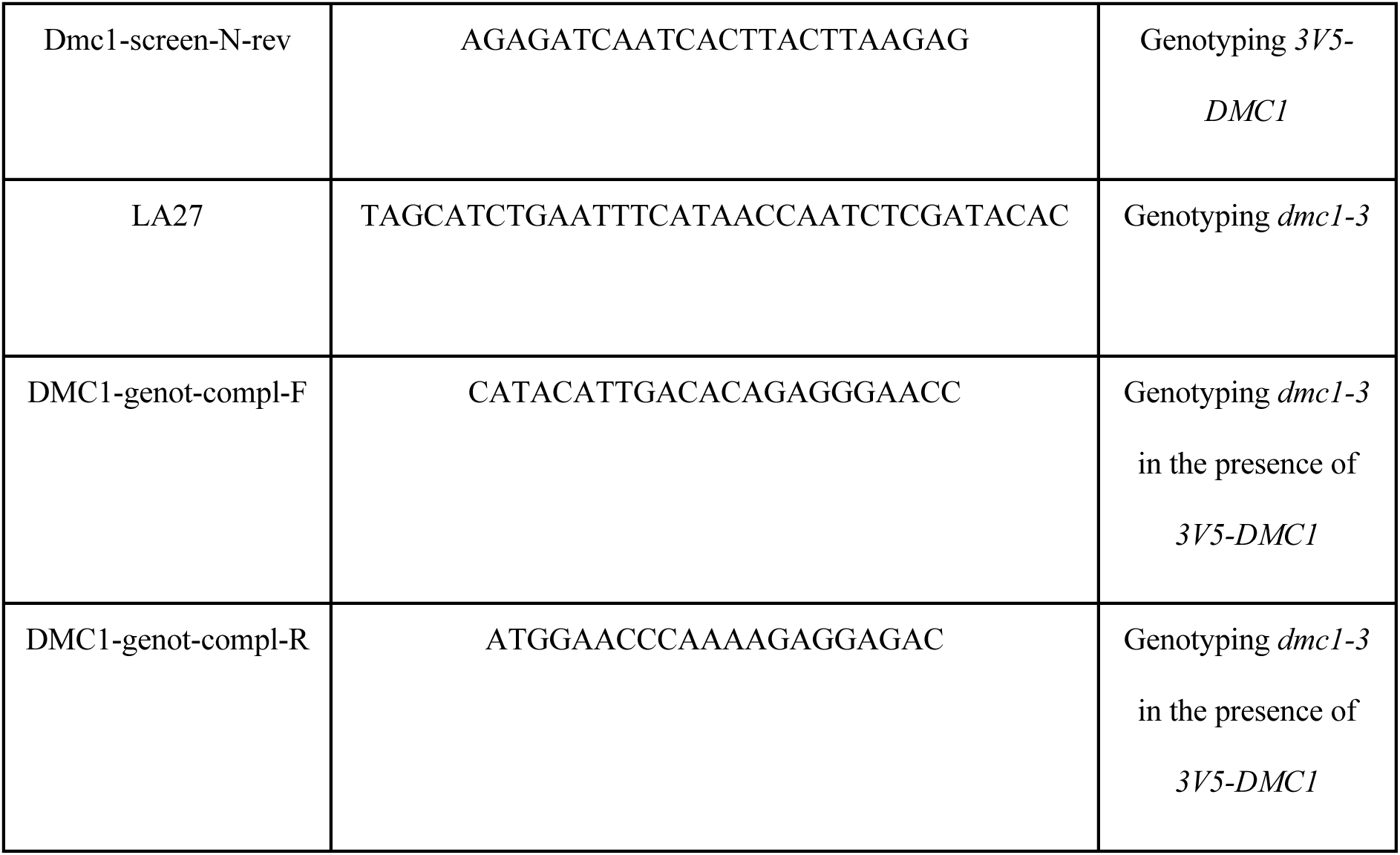
Oligonucleotides. The sequence of oligonucleotides used for *V5-DMC1* construction and genotyping are listed.

**Figure S1.**
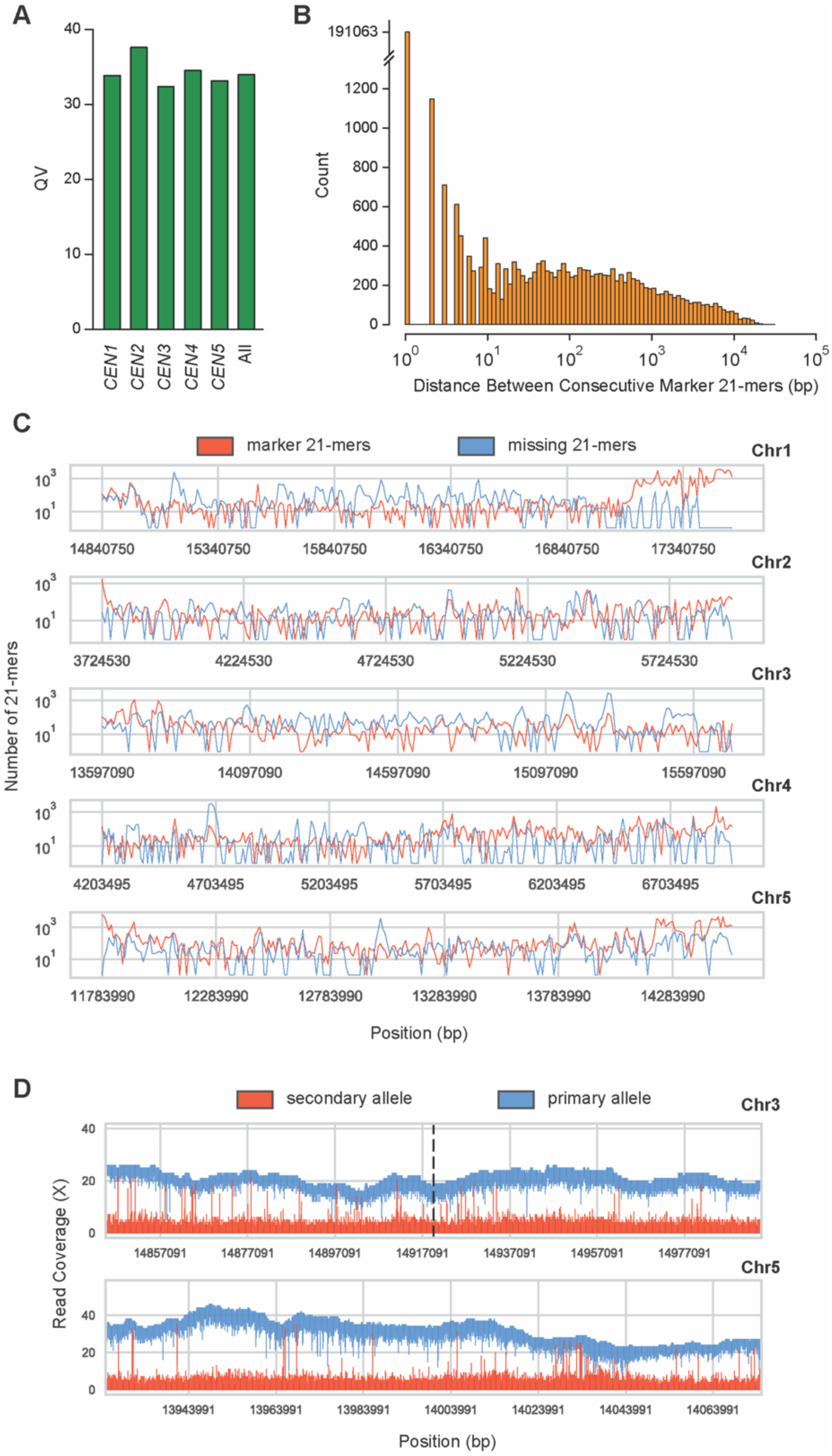
Validation of the Col-CEN centromere assembly. **A.** Assembly consensus quality (QV) scores of the individual and collective (All) centromeres. **B.** Distribution of distances (bp) between consecutive marker 21-mers. **C.** The number of marker (red) and missing (blue) 21-mers in non-overlapping 10 kbp windows across the centromeres. **D.** Primary (blue) and secondary (red) allele ONT coverage for the chromosome 3 *CEN180* inversion region (upper, dotted black line indicates the *CEN180* strand transition), and a chromosome 5 *ATHILA* invaded region (lower).

**Figure S2.**
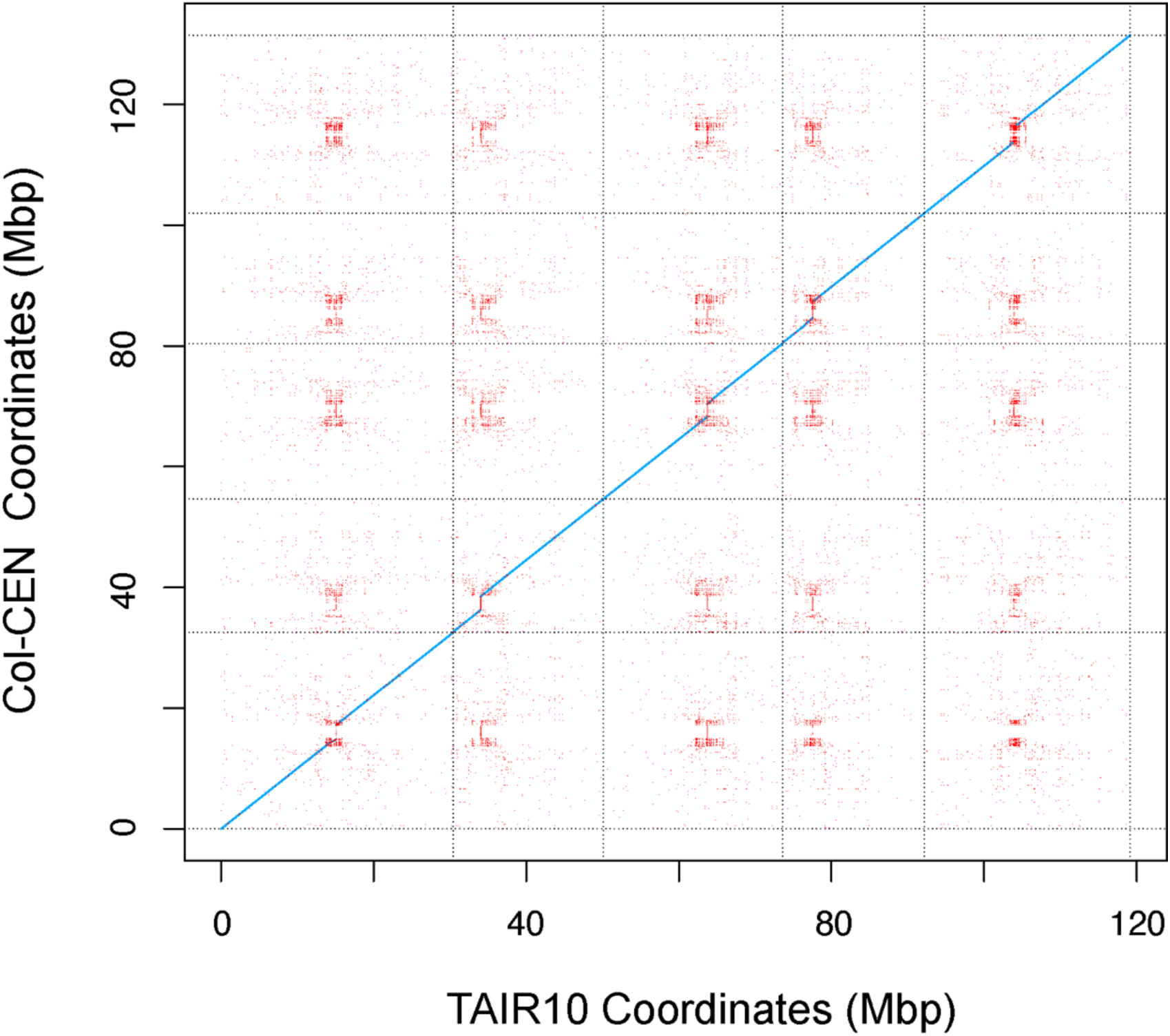
Dotplot sequence similarity comparison of TAIR10 and the Col-CEN genome assembly. A dotplot depicting unique (blue) and repetitive (red) Nucmer alignments (--maxmatch -l 50 -c 250) between TAIR10 and Col-CEN.

**Figure S3.**
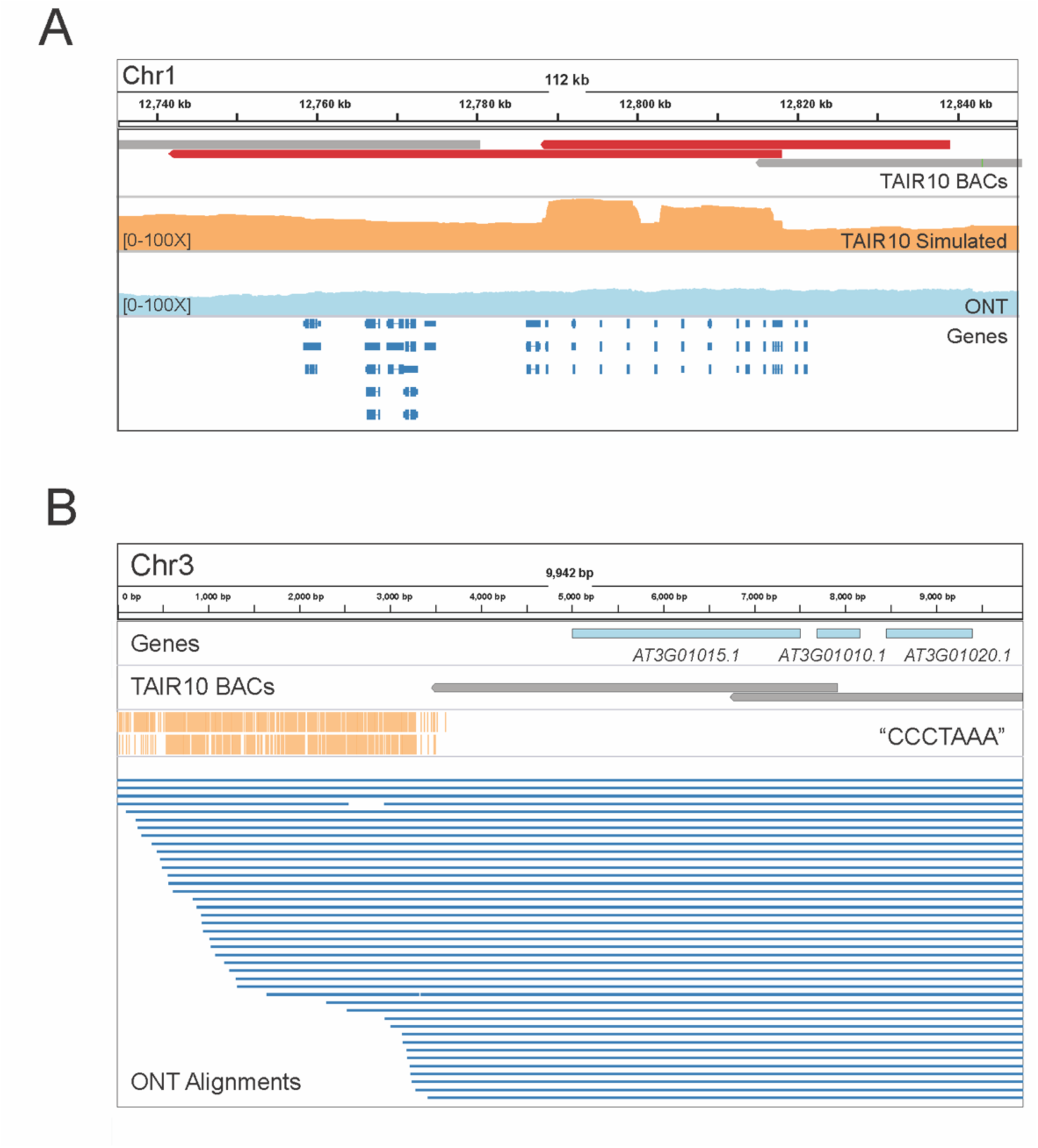
Analysis of thionin gene cluster and telomere assemblies. **A.** A Col-CEN IGV screenshot depicting a thionin gene cluster on chromosome 1. The top track shows TAIR10 BAC contig alignments, with a single BAC contig (red) aligning discordantly. 100 kbp exact WGS reads were simulated from TAIR10 and the coverage of their alignments is shown in orange, followed by ONT alignment coverage (light blue) and gene annotation (dark blue) tracks. As the TAIR10 simulated reads show punctuated coverage increases, but our Nanopore reads do not, this suggests a true biological difference between the sequences. **B.** A Col-CEN IGV screenshot depicting the beginning of chromosome 3, including the assembled telomere.

**Figure S4.**
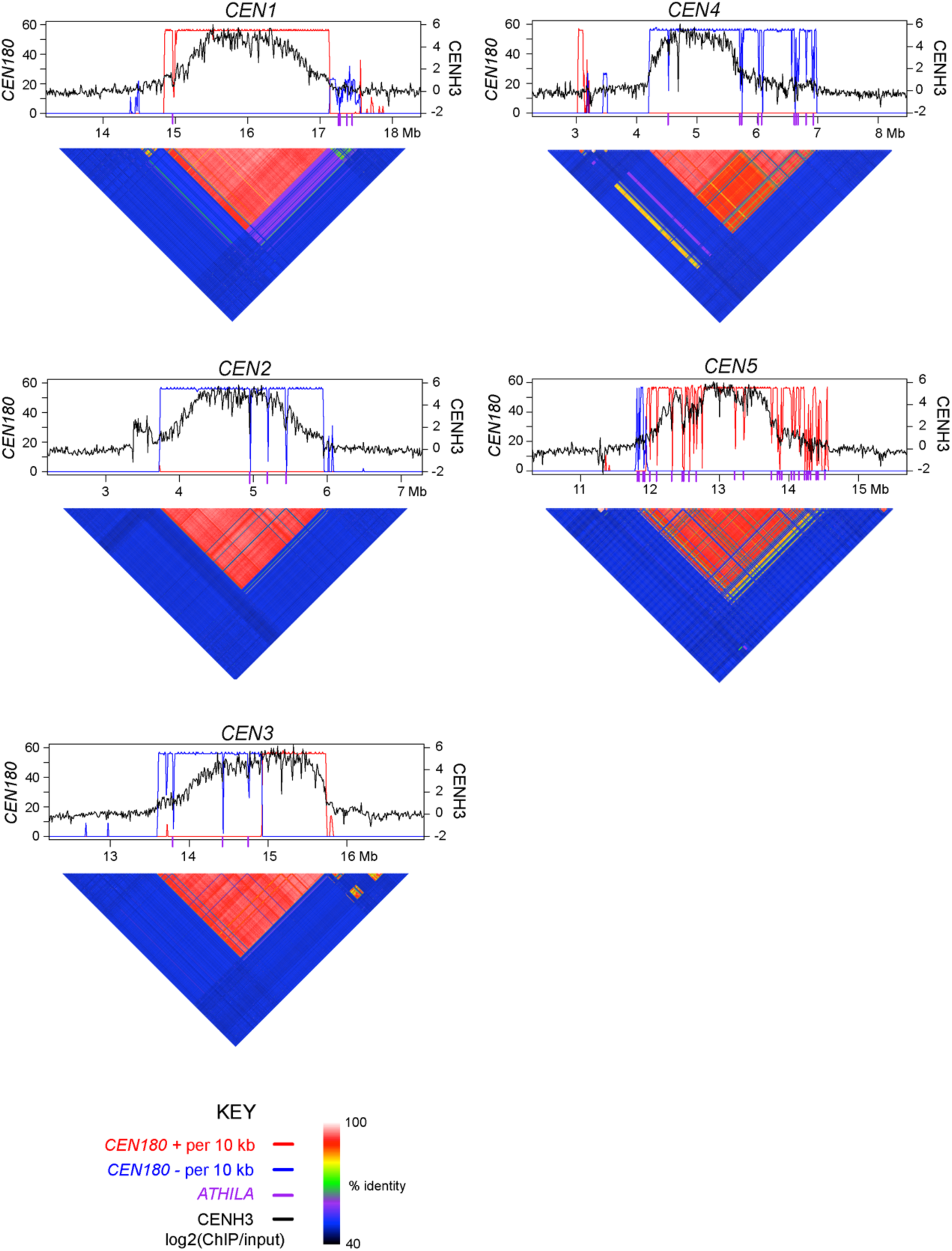
CENH3, *CEN180* and sequence identity across the Arabidopsis centromeres. CENH3 log_2_(ChIP/Input) (black) (*13*), plotted over each centromere. *CEN180* density per 10 kbp is plotted showing forward (red) or reverse (blue) strand orientations. The location of *ATHILA* retrotransposons is indicated by purple ticks on the x axis. Beneath the plot are heatmaps indicating pairwise % identity values between adjacent 5 kbp regions.

**Figure S5.**
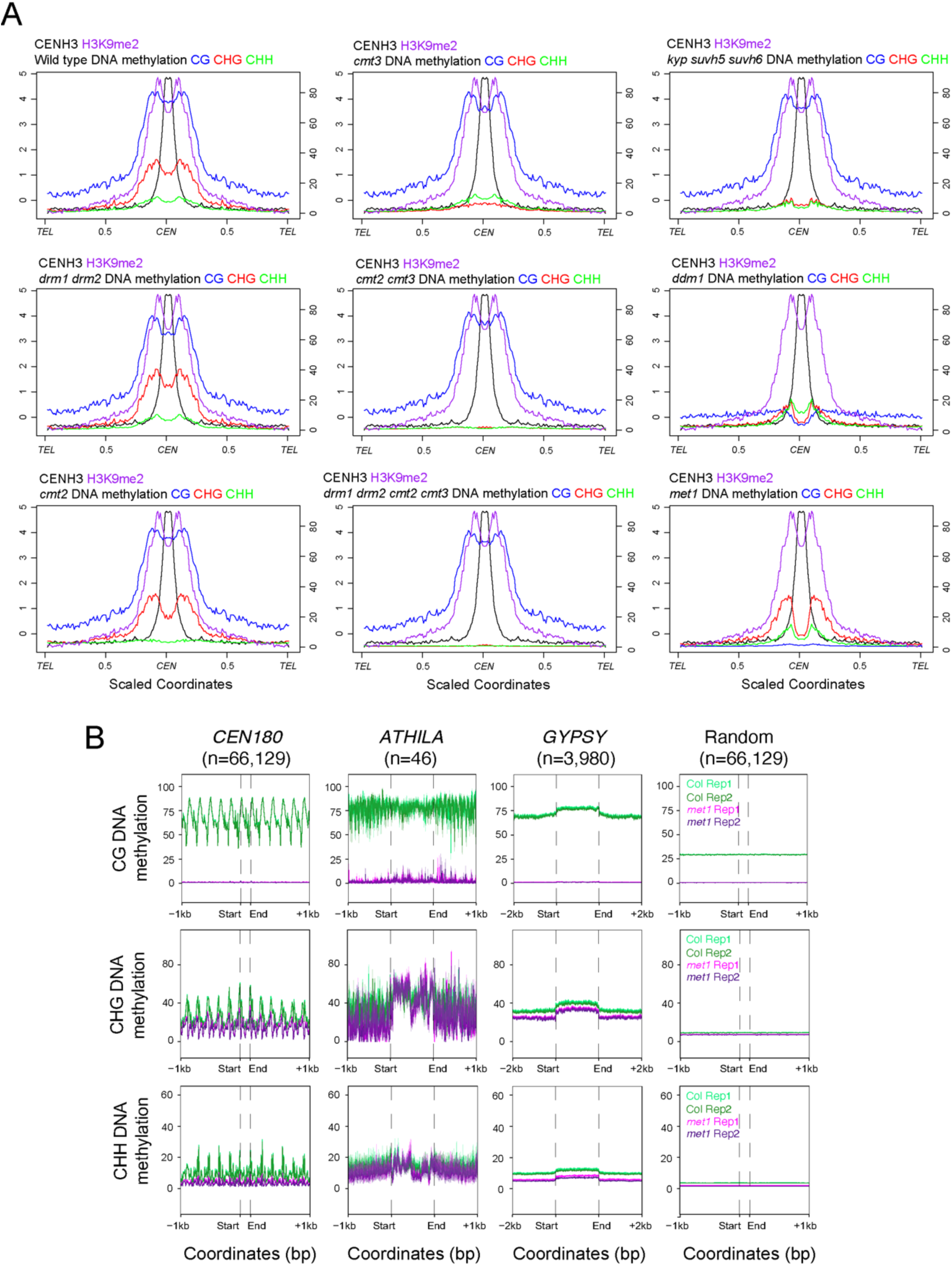
DNA methylation in wild type and CG and non-CG pathway mutants. **A.** Plots of CENH3 (black) and H3K9me2 (purple) log_2_(ChIP/Input) along chromosomes scaled proportionally along the telomere-centromere axes, and centred on the region of maximum CENH3 enrichment (*13, 28*). DNA methylation profiles calculated from BS-seq data are plotted for CG (blue), CHG (red) and CHH (green) sequence contexts in the indicated genotypes (*26*). **B.** CG, CHG and CHH context DNA methylation in wild type (Col-0) or *met1* measured using BS-seq (*31*), over *CEN180* (n=66,129), centromeric *ATHILA* (n=46), *GYPSY* located outside the centromeres (n=3,980) and random positions (n=66,129). Shaded ribbons represent 95% confidence intervals for windowed mean values.

**Figure S6.**
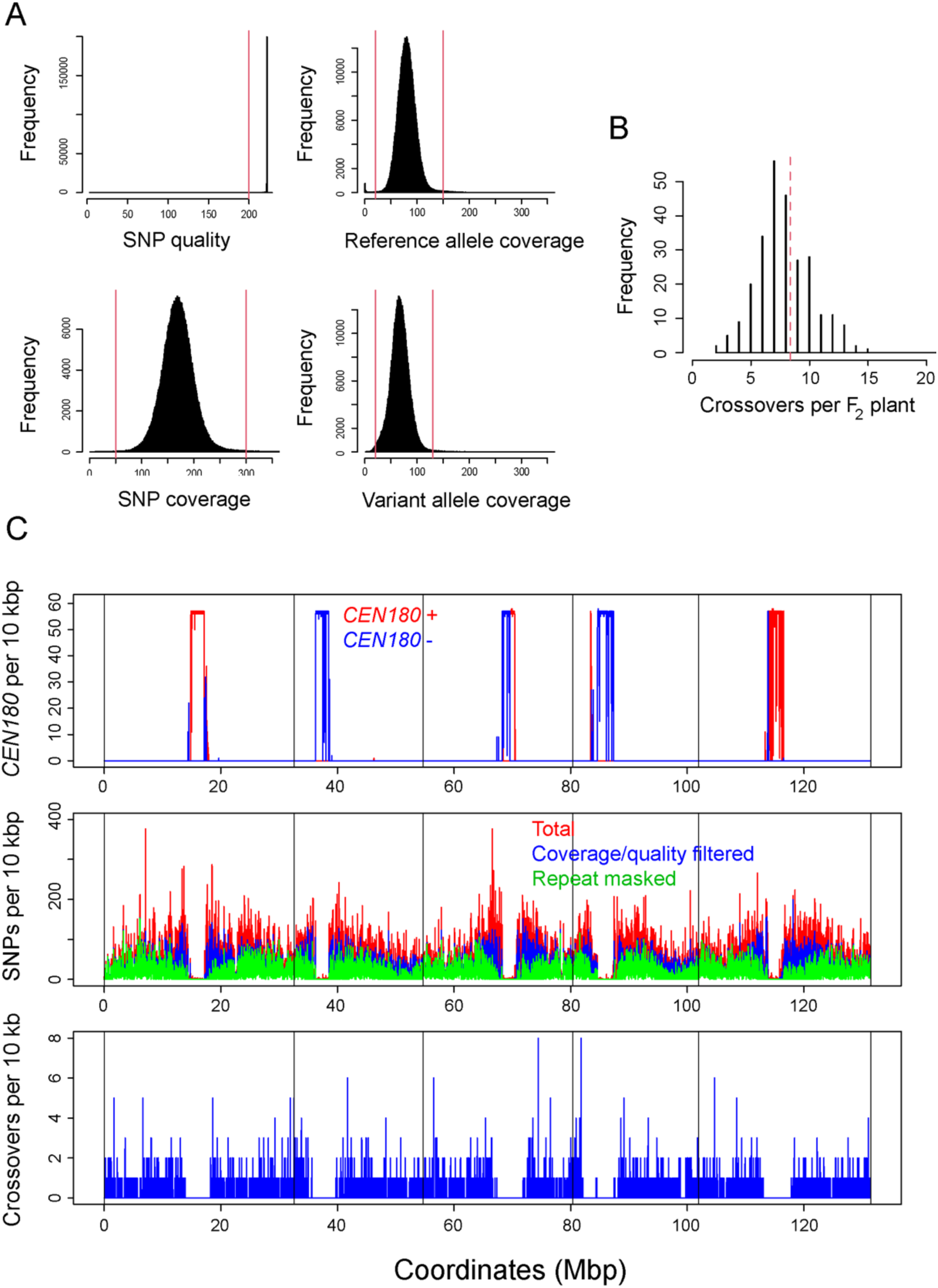
Mapping Col×Ler single nucleotide polymorphisms (SNPs) and crossovers against the Col-0 centromere assembly. **A.** Histograms showing the frequency of qualities, coverage, reference and variant allele coverages for single nucleotide polymorphisms (SNPs) called against the assembly using data from 260 Col×Ler genomic DNA F_2_ sequencing libraries. The red lines indicate thresholds where sites were filtered out of analysis. **B.** Histogram of crossovers mapped against the assembly per Col×Ler F_2_ plant. The red dotted line indicates the mean value. **C.** Plot of the assembly showing *CEN180* satellite density per 10 kbp for forward (red) and reverse (blue) strands (upper). Beneath, the frequency per 10 kbp of total Col×Ler SNPs (red) are plotted, in addition to SNP frequency filtered for quality and coverage values, as in A (blue), and SNPs following repeat-masking (green). The lower plot shows crossovers per 10 kbp (blue) mapped against the assembly.

**Figure S7.**
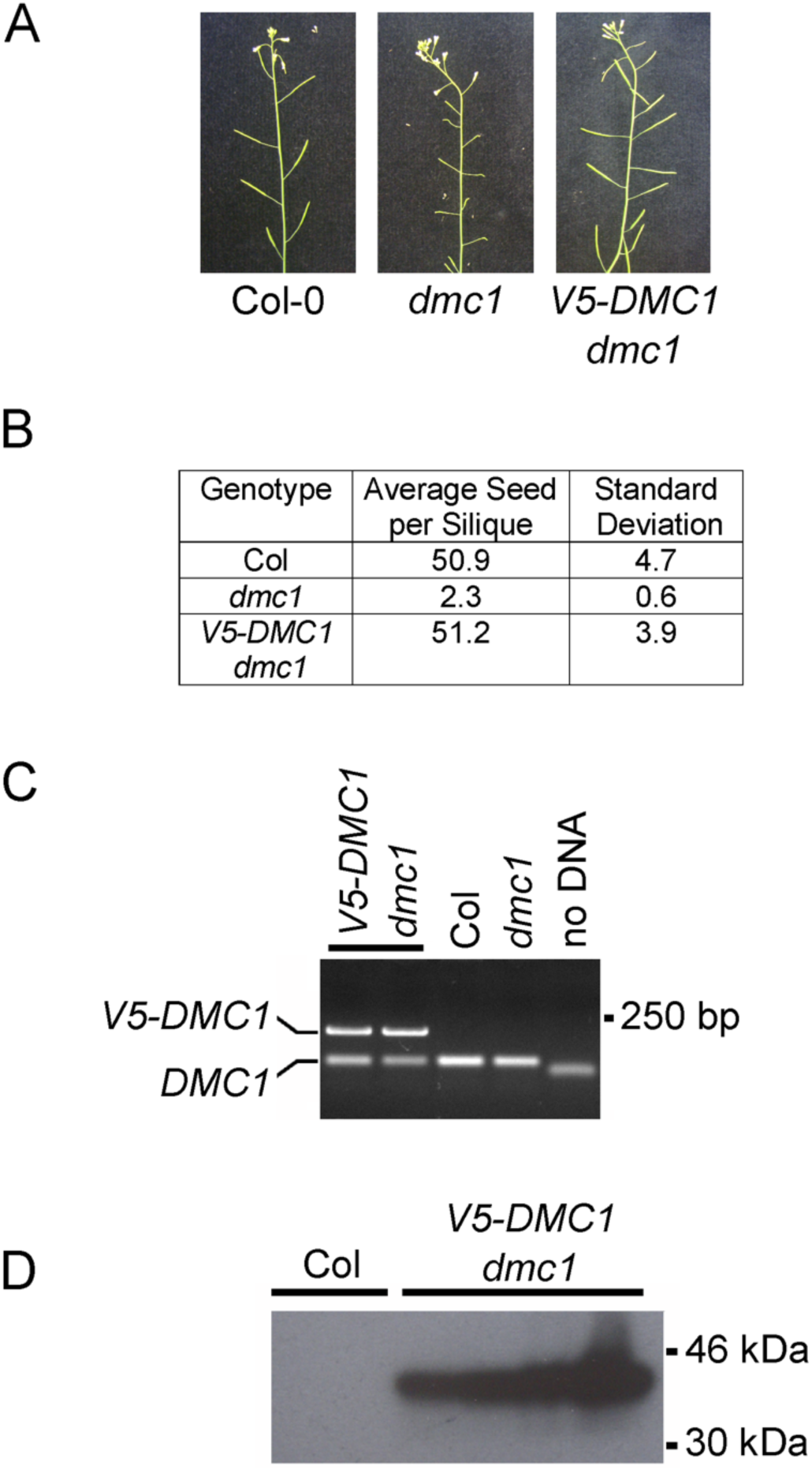
Epitope-tagging and functional complementation of *V5-DMC1*. **A.** Inflorescences of wild type (Col-0), *dmc1-3* and *V5-DMC1 dmc1-3*. Fertility is evident from silique length. **B.** Quantification of seed set per silique in wild type (Col-0), *dmc1-3* and *V5-DMC1 dmc1-3*. **C.** PCR based detection of the N-terminally epitope-tagged *V5-DMC1* transgene, alongside Col-0 and *dmc1-3* null controls. PCR primers flank the *DMC1* ATG translation start site. The expected PCR product sizes are 203 and 74 bp for epitope-tagged and wildtype *DMC1*, respectively. Unincorporated oligonucleotides are seen in ‘no DNA’ control. **D.** α-V5 western blot from Col-0 and *V5-DMC1 dmc1-3* protein extracts from closed flower buds. The expected size of *V5-DMC1* is 41.7 kDa.

**Figure S8.**
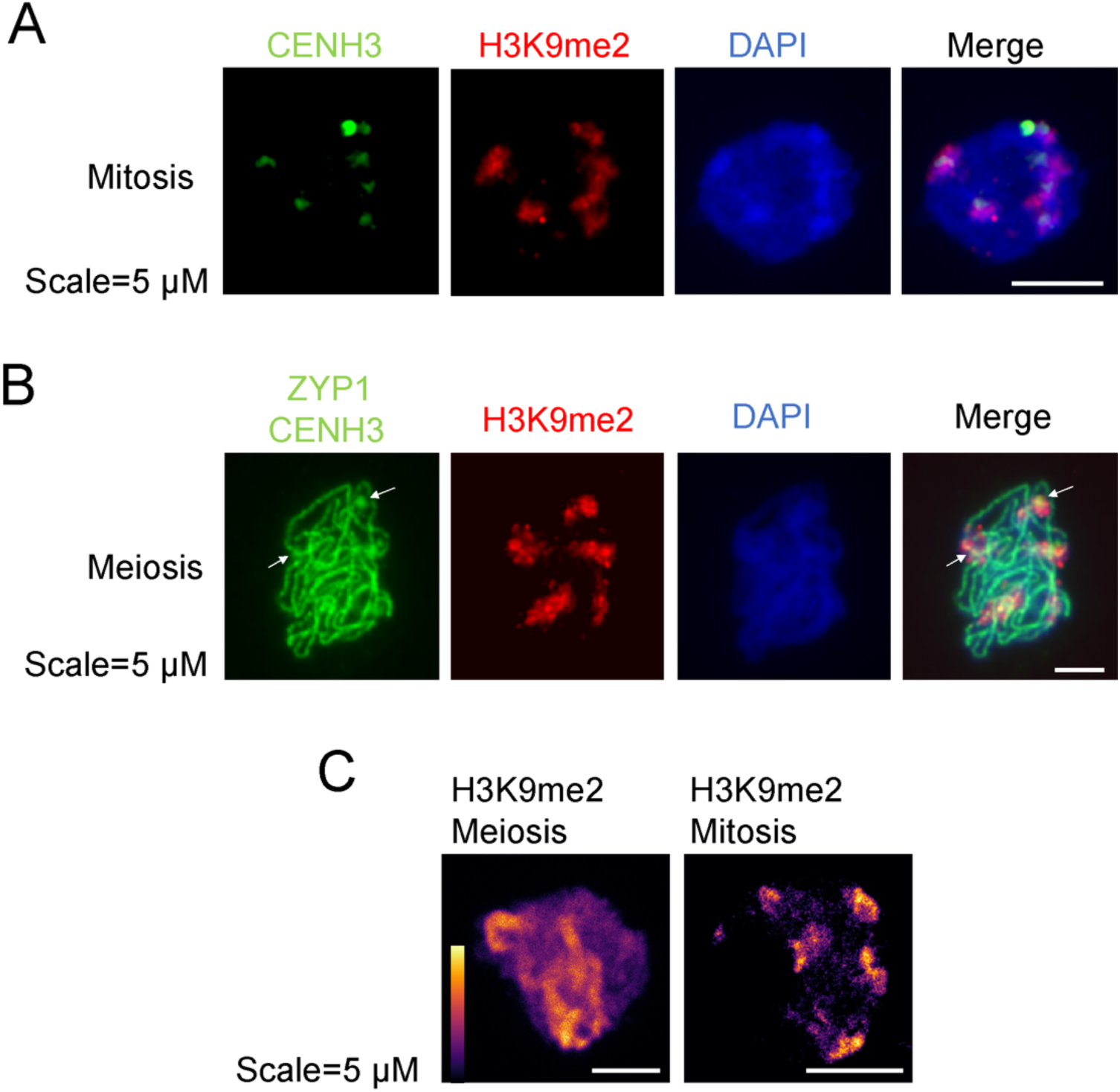
Immunocytological staining of the Arabidopsis centromeres. **A.** Somatic (mitotic) cell immunostained for CENH3 (green), H3K9me2 (red) and stained for DAPI. Scale bar=5 μM. **B.** As for A, but showing an Arabidopsis male meiocyte in pachytene immunostained for CENH3 (green), ZYP1 (green) and H3K9me2 (red), and stained for DAPI (blue). Scale bar=5 μM. **C.** Mitotic and meiotic cells immunostained for H3K9me2 and imaged using STED super resolution microscopy. The colour-scale indicates the intensity of staining, with yellow representing the maximum intensity. Scale bars=5 μM.

